# Structural Analysis of Simultaneous Activation and Inhibition of γ-Secretase Activity in Development of Drugs for Alzheimer’s disease

**DOI:** 10.1101/2020.09.22.307959

**Authors:** Željko M. Svedružić, Katarina Vrbnjak, Manuel Martinović, Vedran Miletić

## Abstract

**Significance:** The majority of drugs that target membrane-embedded protease γ-secretase show unusual biphasic activation-inhibition dose-response in cells, model animals, and humans. Semagacestat and avagacestat are two biphasic-drugs that can facilitate cognitive decline in patients with Alzheimer’s disease. Initial mechanistic studies showed that the biphasic-drugs, and pathogenic mutations, can produce the same type of changes in γ-secretase activity.

**Results:** DAPT, semagacestat LY-411,575, and avagacestat are four drugs that show different binding constants, and biphasic activation-inhibition dose-response curves, for amyloid-β-40 products in SHSY-5 cells. Multiscale molecular dynamics studies showed that all four drugs bind to the most mobile parts in presenilin structure, at different ends of the 29 Å long active site tunnel. Combined results from structure-activity studies, showed that the biphasic dose-response curves are a result of modulation of γ-secretase activity by concurrent binding of multiple drug molecules at each end of the active site tunnel. The drugs activate γ-secretase by forcing the active site tunnel to open, when the rate-limiting step is the tunnel opening, and formation of the enzyme-substrate complex. The drugs inhibit γ-secretase as uncompetitive inhibitors, by binding next to the substrate to dynamic enzyme structures that regulate processive catalysis. The drugs can modulate the production of different amyloid-β catalytic intermediates, by penetrating into the active site tunnel to different depth with different binding affinity. The drugs and pathogenic mutations affect the same dynamic processes in γ-secretase structure.

**Conclusions:** Biphasic-drugs like disease-causing mutations can reduce the catalytic capacity of γ-secretase and facilitate pathogenic changes in amyloid metabolism.

## Introduction

Alzheimer’s disease is a slowly progressing neurodegenerative disorder with a fatal outcome *(1)*. Alzheimer s disease is also the major challenge for the pharmaceutical industry today, as it stands out ahead of the malignant diseases, as the biggest financial burden for the health care providers in developed countries *(1, 2)*.

The great majority of the potential drugs for Alzheimer’s disease have targeted the metabolism of the C-terminal part of amyloid precursor protein (C99-APP) *(2-5)*. The target enzyme was the membrane-embedded aspartic protease: γ-secretase *(2-4)*. A large number of repeated failures in the last twenty years have led to numerous proposals that γ-secretase might not be a good therapeutic target *(2-5)*. An alternative, less frequently considered explanation, is that a very few of the failed drug-design strategies, have taken into account the complexity of the catalytic mechanism of γ-secretase *(2, 4-7)*.

Activation and inhibition of γ-secretase activity that can be observed in biphasic doseresponse to drugs, are good examples of complex enzymatic mechanism that was not adequately recognized in drug-development studies *(2, 6, 8-14)*. Biphasic dose-response can be observed in cell-cultures *(8, 10, 12, 15)*, in model animals *(8, 14)*, and in patient’s plasma in clinical trials *(9)*. The biphasic dose-response can be observed only in studies that use physiological sub-saturating substrate concentrations, and a full concertation range for the drugs *(7-10)*. It is very likely, that all types of drugs that target presenilin subunit of γ-secretase, can produce the biphasic dose-response when Aβ metabolism is at the physiological level *(6, 8, 9, 12)*.

The great majority of drug-screening and optimization studies used unphysiological high saturation of γ-secretase with its substrate and a limited concentration range for the drugs. Such an approach can be a good time and money-saving strategy in the early screening process *(16-18)*. However, the “fast-and-cheap” approach is also the main reason why the majority of the preclinical studies were misleading, or had poor relevance to the clinical studies *(2, 4, 6, 8, 9, 13, 14)*. γ-Secretase is far from saturation with its substrate in physiological conditions in cells *(7, 8, 10, 14)*, just as the majority of other enzymes *(16, 19, 20)*. Sub saturated enzymes are a fundamental physiological mechanism, that can assure the fastest, and best-controlled responses to metabolic changes *(16, 19)*. Measurements that had poorly defined saturation of γ-secretase with its substrate, can explain why so many studies have frequently reported irreproducible results on selectivity between Notch and APP substrates, or on modulation of Aβ production *(2, 4, 13)*. Reproducible experiments depend on well-defined Kӎ values for different substrates and Aβ products *(12, 16, 17, 21, 22)*.

The initial mechanistic studies showed that the biphasic dose-response drugs can produce the same type of changes in γ-secretase activity as the disease-causing FAD mutations *(7, 23)*. Both, FAD mutation and the biphasic-drugs, can cause accumulation of the longer and more hydrophobic Aβ products *(6, 12, 21, 24, 25)*. At low physiological substrate concentrations, the biphasic-drugs can increase the saturation of γ-secretase with its substrate like the Swedish mutation in its APP substrate *(10)*. At high substrate concentrations, the biphasic-drugs act as uncompetitive inhibitors and show a decrease in the maximal turnover rates like the FAD mutations in presenilin 1 *(7, 10, 21, 24)*. In essence, FAD mutations and biphasic-drugs have the same effect, a decrease in γ-secretase capacity to process its substrates *(7)*. A decrease in the catalytic capacity of γ-secretase can produce pathogenic changes in Aβ products with the wild type γ-secretase *(7, 21, 22, 26)*. Dose-dependent cognitive decline was observed in clinical trials with biphasic-drug avagacestat *(9, 27)* and probably with semagacestat *(6, 15, 28-30)*.

Here we present combined structure and activity studies of γ-secretase, to describe the biphasic dose-response in the presence of four best-known drugs. Semagacestat and avagacestat were chosen as two biphasic-drugs that can facilitate cognitive decline in patients in clinical trials *(6, 9, 13-15, 27, 30)*. DAPT is chosen as a biphasic-drug that has been most frequently used in mechanistic studies of γ-secretase *(10, 12, 25, 31-33)*. LY-411,575 is chosen as a biphasic-drug with one of the most potent affinities for γ-secretase *(34)*. We found that the multiple drug molecules can bind simultaneously to the different flexible parts on the presenilin subunit of γ-secretase. The drugs bound to different sites can affect every catalytic step: from the enzyme-substrate recognition to the processive cleavages of Aβ products. We also found that the drugs and FAD mutations target the same dynamic parts in the presenilin structure.

## Results

### Biphasic activation-inhibition dose-response curves for DAPT, semagacestat, LY-411,575 and avagacestat (Figs. 1-2)

We measured biphasic dose-response in Aβ 1-40 production in SHSY-5 cells *(8, 10, 14, 15)*. The biphasic dose-response depends on the extent of the γ-secretase saturation with its substrate and the assay design *(7, 8, 10)*. Different drugs can be quantitatively compared only if the drugs were incubated under identical conditions in identical cells. Here presented measurements stand out as the measurements that have taken maximal efforts to make identical assay conditions for each of the four drugs (Fig. 1). One batch of cells was split into four identical batches for parallel treatment with the four drugs in exactly the same conditions. The cells were incubated with each drug overnight, and the Aβ 1-40 production was measured in parallel using identical protocols.

**Figure 1.**
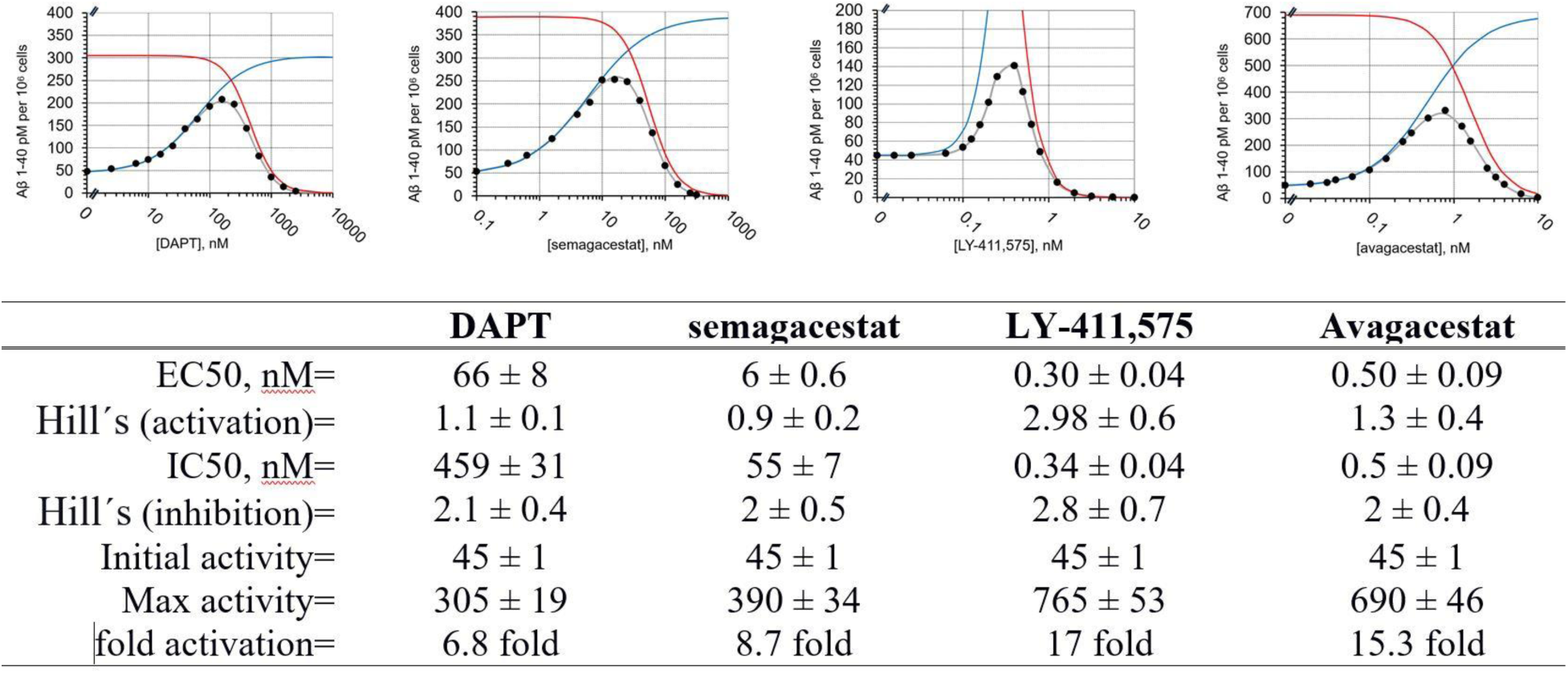
Dose-response curves for Aβ 1-40 production in SHSY cells were measured in the presence of DAPT, semagacestat, LY-411,575, and avagacestat. Aβ 1-40 production in SHSY-5 cells shows biphasic activation-inhibition dose-response curves with all four drugs. Different parameters that describe the biphasic dose-response curves were calculated and listed in the table (eqn. 1) *(10)*. The gray lines represent the best-fit curve to experimental values that are represented by black dots. The blue and red lines represent calculated activation and inhibition events if the two events can be separated. Activation constants (EC50) and the inhibition constant (IC50) represent the affinity for each binding event. The Hill’s coefficients represent the stoichiometry of interaction, and/or possible cooperative processes in the binding events. “Max activity” parameter represents the maximal possible activation if there is no competing inhibition. This parameter roughly correlates with the ability of drugs to facilitate enzyme-substrate interactions *(10)*. The initial activity is the same for all four drugs because all measurements used the same batch of cells.

Aβ 1-40 production in SHSY-5 cells treated with DAPT, semagacestat, LY-411, 575, and avagacestat show biphasic activation-inhibition dose-response curves with clear differences between the drugs in activation and inhibition constants and Hill’s coefficients (Fig. 1). Thus, the structure-activity analysis of these molecules can give insights into the mechanism behind biphasic responses (Fig. 2). Full numerical analysis of biphasic dose-response curves depends on five best-fit parameters (Fig. 1) *(10)*. The parameters can be calculated with satisfactory accuracy if the experiments have an optimal selection of concertation range for each drug (Fig. 1) *(7, 10)*.

**Figure 2.**
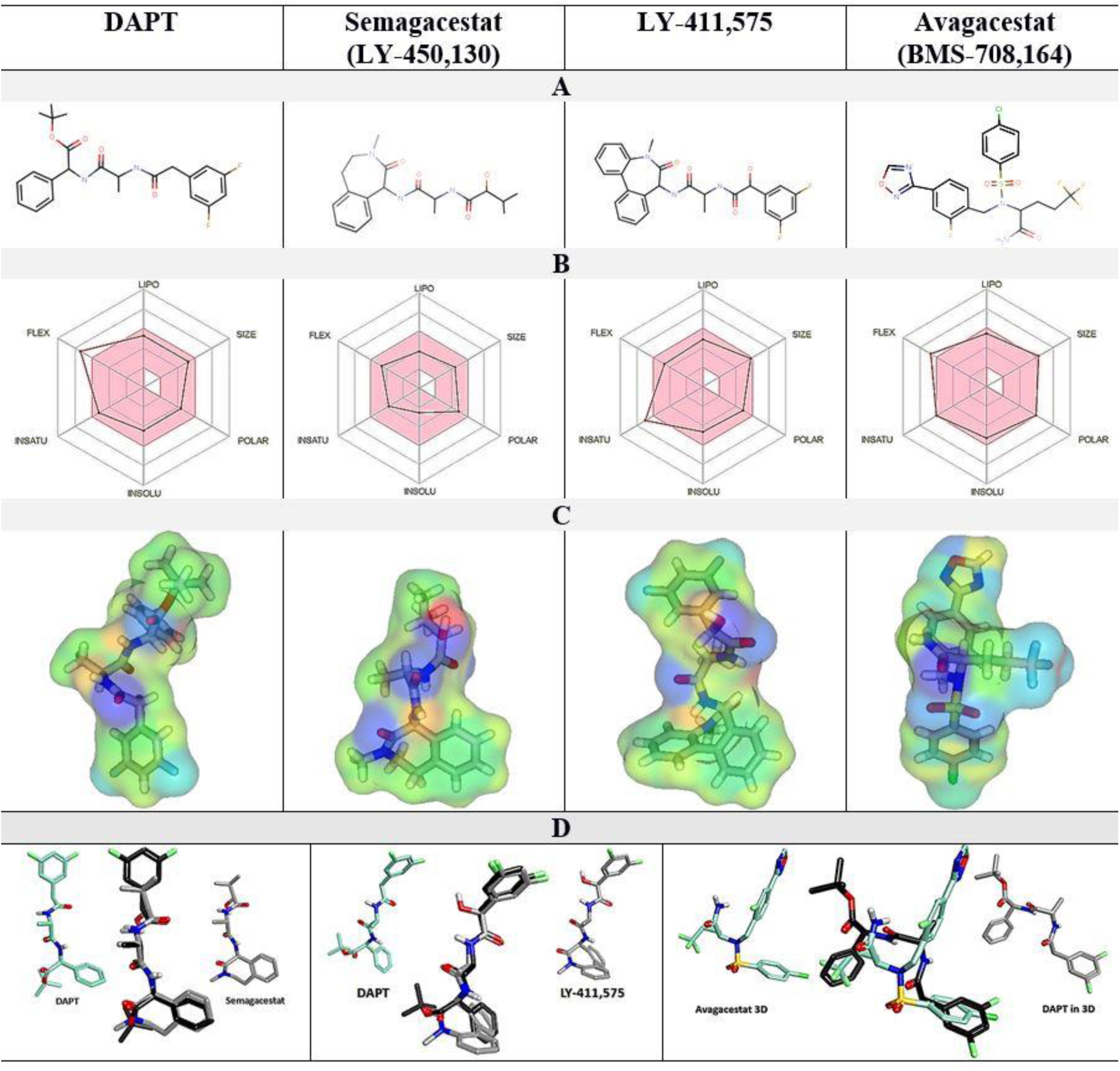
(A-D). Comparative analysis of physicochemical properties of DAPT, semagacestat, LY-411,575, and avagacestat structures. (**A**) 2D structures. (**B**) a radar diagram of overlap with the Lipinski rules: FLEX flexibility, INSAT relative share of sp^3^ carbon atoms, INSOLU LogP values, POLAR polar surface area, SIZE molecular mass, LIPO hydrophobic surface area. The pink area represents the optimal values, the superimposed lines represent the values specific for each compound. (**C**) electron densities are mapped on molecular surfaces and colored to highlight the surface properties: green hydrophobic, blue H-bond donor, red H-bond acceptor, yellow polar. (**D**) overlap of 3D molecular structures with DAPT as the reference molecule.

Five parameters can define the biphasic activation-inhibition profiles: activation (EC50) and inhibition (IC50) constants represent the binding affinity for each binding event *(10, 16, 17)*. Hill’s coefficients represent the stoichiometry of interaction, and/or possible cooperative processes in the two binding events. “*Max activity*” parameter represents the maximal possible activation if there is no competing inhibition present. This parameter roughly correlates with the capacity of each drug to facilitate enzyme-substrate interaction *(10)*. The initial activity parameter is the same for all four drugs, because all measurements used the same batch of cells (Fig. 1).

We find that the activation and inhibition curves always overlap, and that compounds that show low EC50 values for activation also show low IC50 for inhibition (Fig. 1). These results indicate concurrent binding at the activation and inhibition sites *(10, 11, 17)*. The biphasic profiles show Hill’s coefficients that are higher than one (Fig. 1), which indicates the simultaneous binding of multiple drug molecules to γ-secretase and possible cooperativity *(10, 17)*. The inhibition phase shows higher Hill’s coefficients than the activation phase, which could indicate that binding at the activation site can facilitate binding at the inhibition site. There is no correlation between IC50 and EC50 constants and Hill’s coefficients (Fig. 1).

DAPT, semagacestat, LY-411,575 show quantitative differences in biphasic dose-response curves (Fig. 1). DAPT, semagacestat, LY-411,575 show a very close overlap in 3D structures but have different physicochemical properties (Fig. 2). The compounds have the same peptide backbone but differ in the molecular volume and flexibility of aromatic rings at the C and N terminus (Fig. 2). LY-411,575 has the lowest EC50 values for activation and IC50 values for inhibition, the highest possible activation levels, and the highest Hill’s coefficients (Fig. 1). DAPT shows the highest EC50 and IC50 constants with the lower Hill’s coefficient and lower maximal possible activation (Fig. 1). The biphasic parameters measured with semagacestat are between DAPT and LY-411,575 values.

Avagacestat and LY-411,575 have similar EC50 and IC50 and notable differences in the Hill’s coefficients (Fig. 1). Avagacestat stands out, as a different structure, which cannot overlap with the other three molecules in 3D alignments of molecular structures (Fig. 2).

All four compounds have more than 5 hydrogen bond donor and acceptor sites, that are surrounded by large hydrophobic surfaces at the terminal ends of the molecules (Fig. 2). The most notable difference between the four molecules is in the molecular volume (Fig. 2), which indicates that the four compounds can bind with different affinity to the different size cavities on the γ-secretase surface.

### Multiscale molecular dynamics studies of γ-secretase structure in different steps in the catalytic cycle (Fig. 3)

The flexible parts in the γ-secretase structure can be described using residue-based coarse-grained protocols (Fig. 3, *(35)*). These protocols can routinely depict flexible protein parts in as much as 20 microseconds of molecular events (Fig. 3) *(35)*. We are specifically interested in enzyme-substrate interaction (Fig. 3) *(36)*, and in the processing of different Aβ catalytic intermediates (Fig. 3) *(12, 36)*. Those complexes are most frequently targeted in different drug development strategies *(8, 10, 12)*.

**Figure 3.**
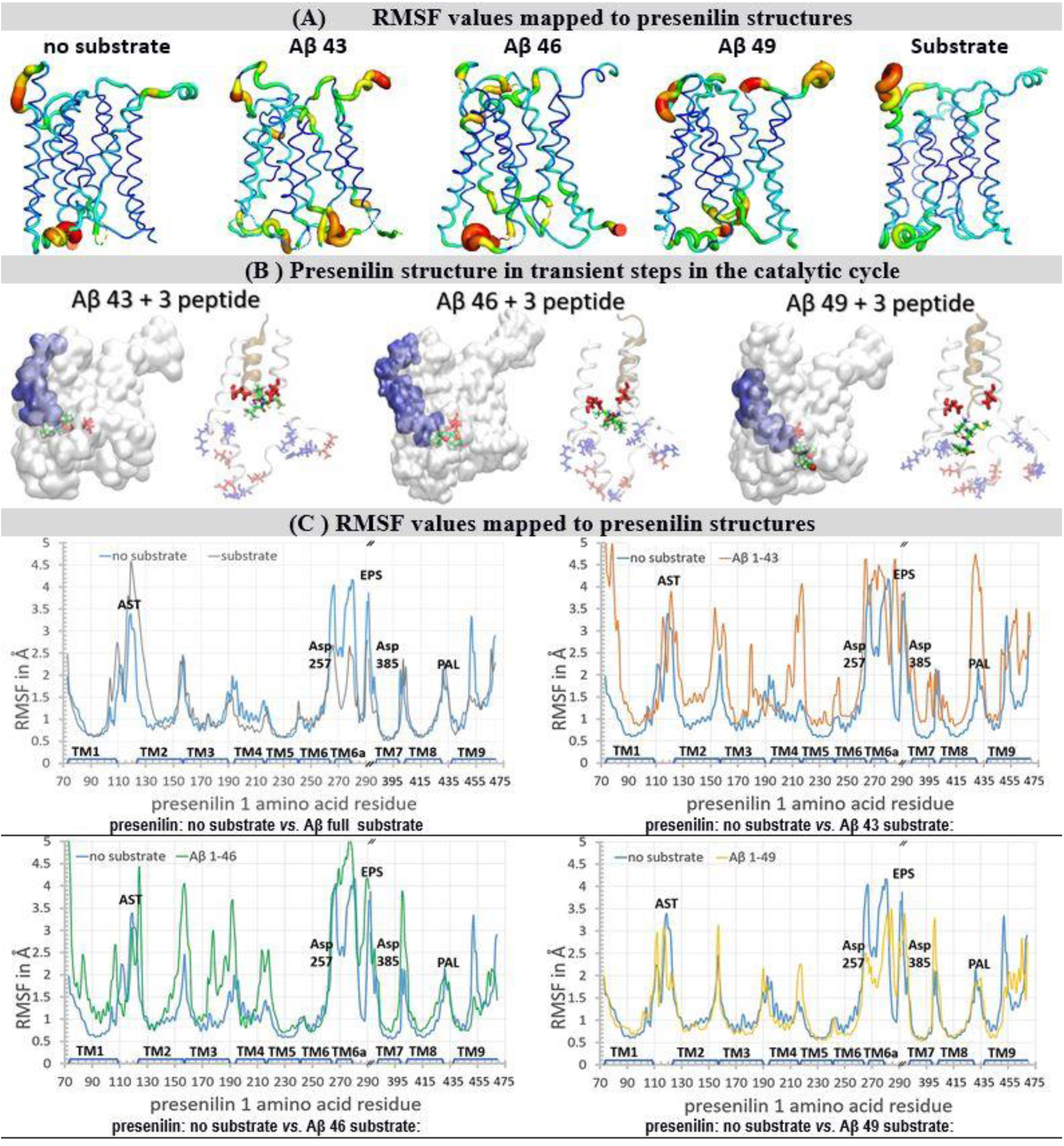
(<u>A</u>-<u>C</u>). Residue-based coarse-grained molecular dynamics studies of γ-secretase complex. We used cryo-EM coordinates (PDB:6IYC) *(36)* to describe how substrate and Aβ catalytic intermediates can affect the dynamic conformational changes in the γ-secretase complex. Described conformation changes represent between 10 to 20 µsec of molecular events *(35)*. The results are depicted visually (A-B), and quantitatively (C), with the focus on presenilin 1 structures. **(<u>A</u>)** Principal component analysis showed that the substrate and Aβ catalytic intermediates can affect the mobility of presenilin 1 structure at the secondary structure level *(38, 39)*. The relative differences in mobility are depicted by shape and color: thin blue lines represent the lowest mobility, green and yellow lines represent intermediate mobility, while tick red lines represent the highest mobility. **(<u>B</u>)** The presenilin structures with Aβ catalytic intermediates show that different complexes are unstable to a different extent, because the structures with partial occupied active site tunnel represent a mixture between the open and closed structures *(36, 37)*. The structures are shown as a white transparent surface to make the structures underneath the surface visible. Different Aβ catalytic intermediates in the active site tunnel are depicted as blue surfaces, active site Asp 257 and Asp 385 are shown as red licorice. The white ribbon models depict amino acids 240 to 394 in presenilin structures, while gold ribbons depict different Aβ catalytic intermediates. The positively charged amino acids are shown as blue licorice, negatively charged amino acids are shown as red licorice, including the active site Asp 257 and Asp 385. Tripeptide by-products of processive catalysis are shown as green models *(12, 25, 36)*. **(<u>C</u>)**RMSF graphs give a quantitative description of structural changes at the level of each amino acid *(38, 39)*. The biggest variability in RMSF values is observed in the structural parts that drive the enzyme-substrate interaction and the processive catalysis *(12, 25, 36)*. The key structural elements are mapped on the graph *(36)*: TM1-TM9 <u>T</u>rans <u>M</u>embrane helix 1 to 9; AST membrane-embedded opening of the <u>A</u>ctive <u>S</u>ite <u>T</u>unnel; EPS <u>E</u>ndo-<u>P</u>roteolytic <u>S</u>ite; PAL motif (<u>P</u>ro 435, <u>A</u>la 434, <u>L</u>eu 433) *(40)*; Asp 257 and Asp 385 in the active site.

Substrate binding and Aβ catalytic intermediates lead to increase in width and mobility in the presenilin structure at its cytosolic end (Fig. 3A) *(36)*. This part is exceptionally rich in charged amino acids that form dynamic salt-bridges (Fig. 3 B, *(36)*, supplement Fig. 1). The presenilin structures with Aβ catalytic intermediates show that different complexes are unstable to a different extent, because the structures with partial occupied active site tunnel represent a mixture between the open and closed structures *(36, 37)*.

The highest mobility is observed with Aβ 43 and Aβ 46 catalytic intermediates (Fig. 3A and 3C). These highly mobile structures represent an unstable transient protein-substrate complex in processive catalysis (Fig. 3B, supp. Fig. 1). With Aβ 43 and the shorter Aβ products, the C-terminal end of the substrate forms repulsive interactions with the Asp 257 and Asp 385 in the active site (Fig. 3B). This results in a strong negative field that attracts compensating interactions from the positively charged amino acids, most notably Lys 380 and Lys 267 (Figs. 3B, and supp. Fig. 1A). The Aβ 46 and longer Aβ products can reach the presenilin interior beyond the active site aspartates (Fig. 3B). There the negatively charged C-terminal makes dynamic slat bridges with different Lys and Arg residues at the cytosolic end of the tunnel (Fig. 3B, supp. Fig. 1). The presenilin in complex with Aβ 49 does not depend on such large conformational changes to engage in catalysis and shows lower mobility (Fig. 3 B-C). The presenilin structure with a full substrate has lower mobility than the no-substrate structure (Fig. 3C). The structure with full substrate depends on the local conformational changes to engage in the catalysis *(36)*, the no-substrate structure has to support much larger conformational changes to engage in catalysis *(36)*.

When Aβ substrates of different lengths are bound to γ-secretase the active site tunnel is predominately closed at its cytosolic end (Fig. 3B and supp. Fig. 1B). The active site tunnel can instantly close in molecular dynamics studies when the substrate is removed (supplement Fig. 1). The closed structure is necessary to prevent leaking of the ions across the membrane *(36)*. The tunnel closing is driven by the dynamic hydrophobic interactions between the amino acids that form the tunnel walls and the flexible protein loops at each end of the tunnel (Supp. Figs. 1-3, *(36)*). The highly dynamic conformational changes drive the different ends of the closed substrate tunnel open to a different extent (Fig. 3A). In the longest molecular dynamics simulations, that can depict 20 microseconds of molecular events, in only about 0.4% of the simulation time two ends of the substrate tunnel can be connected. These results indicate that the substrate-analogs or mechanism-based inhibitors can penetrate to the active site aspartates only in a series of conformational changes.

RMSF graph can trace the changes in the molecular dynamics at the level of individual amino acids (Fig. 3) *(38, 39)*. RMSF graphs showed that the highest mobility sites are localized to the flexible loops at the different ends of the active site tunnel and most notably at the cytosolic end of the presenilin structure (Fig. 3). The RMSF graphs showed that high mobility sites overlap with the known drug-binding site and the hotspots for disease-causing mutations *(37, 40-45)* (https://www.alzforum.org/mutations). The active site aspartates are part of helix structures and show low mobility deep in the protein interior, however, they are directly adjacent to the high mobility sites (Fig. 3C) *(38, 39)*.

### Search for drug binding sites using molecular docking studies (Fig. 4) (46)

Docking studies used γ-secretase structures with and without the substrate *(36, 37)*. Such an approach can describe competition or cooperation between the substrate and the drugs in binding to the enzyme *(8, 10)*. γ-Secretase is predominately in the substrate-free form in physiological conditions in cells *(10)*.

**Figure 4.**
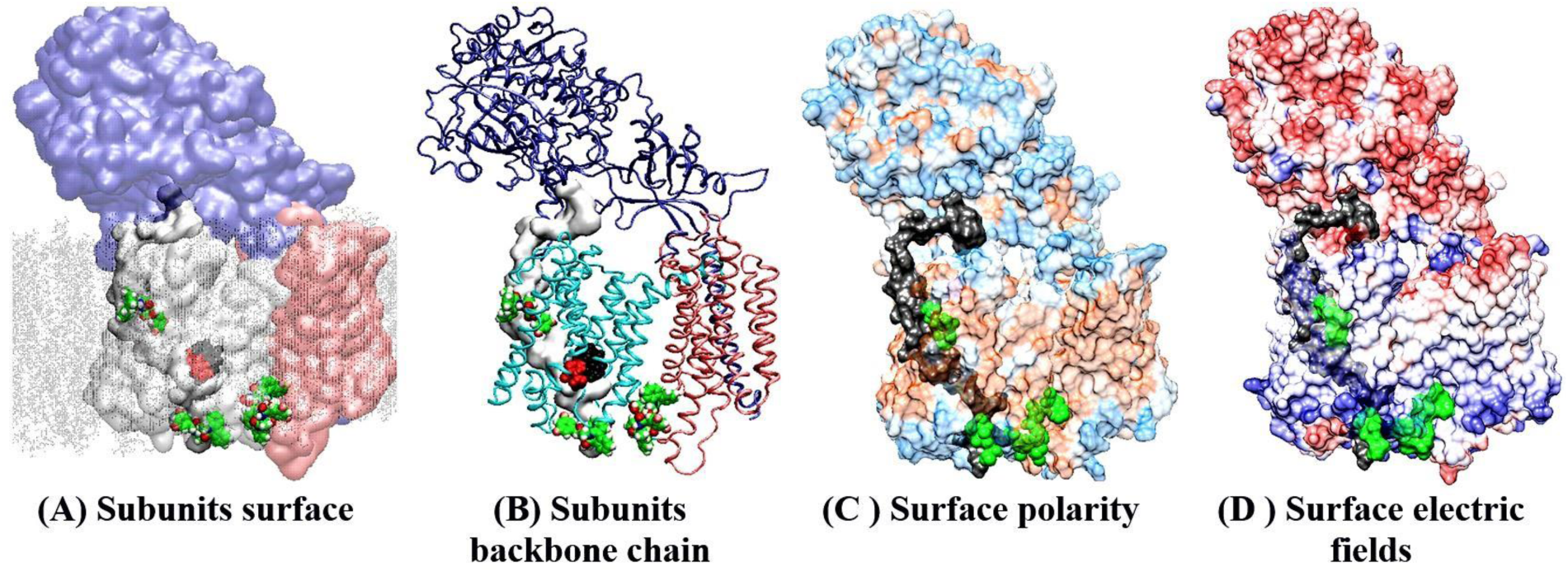
(A-D). Docking sites for semagacestat on the γ-secretase structure in complex with its substrate (PDB:6IYC) *(36)*. Molecular docking studies showed that as much as four semagacestat molecules (green) can bind to γ-secretase simultaneously in presence of the substrate. One drug molecule binds at each end of the active site tunnel, two drug molecules bind in the gap between presenilin and Aph1 subunit. Transparent protein surfaces are used to show the structures underneath the surface. Four different presentations show the position of the drug-binding sites relative to the main structural elements in the γ-secretase complex. (**<u>A</u>**-**<u>B</u>**) Different subunits are shown in different colors, nicastrin cyan, presenilin white or light blue, Aph1 pink, substrate gray, while the membrane is shown as dots. The drugs bind to presenilin sites inside and outside of the membrane. Buried underneath the protein surface is the substrate depicted as a gray surface, the active site Asp 257 and Asp 385 (red), and the adjacent PAL motif (black, Pro 435, Ala 434, Leu 433) *(40)*. (**<u>C</u>**) The protein surface is colored based on its polarity: blue polar, brown hydrophobic, white amphiphilic. The substrate is shown as a black surface, that can also mark the position of active site tunnel (**<u>D</u>**) APBS analysis of electric fields on the protein surface: blue positive, red negative, white neutral *(70)*. The electric fields show that γ-secretase is a polarized molecule. The negative field dominates on the nicastrin side of the membrane, while the positive field dominates the cytosolic site. Thus, the positive N-terminal of the substrate is matching the negative field on nicastrin, and the negative C-terminal on the nascent Aβ catalytic intermediates is matching the positive field at the cytosolic side of the protein. Dynamic electric fields can be a crucial part of enzyme-substrate recognition and processive cleavages of the Aβ catalytic intermediates *(12, 25, 36)*.

The presented docking studies have identified the same binding sites as the earlier studies which used a variety of experimental techniques (Fig. 4) *(37, 42-45)*. The crucial new insights from the presented docking studies is that presenilin can bind multiple drug molecules simultaneously even in the presence of the substrate (Fig. 4). The drugs and the substrate bind to the same sites (Fig. 4), which implicates that some type of competitive or cooperative mechanisms can exist just as indicated in the binding studies (Fig. 1) *(8, 10)*.

The drugs bind to the most mobile parts in the presenilin structure (Figs. 3-4), which regulate the opening of the 29 Å long active site tunnel *(36, 37)*, and thus, must have effects on substrate binding and processive catalysis *(12, 25)*. The binding sites for the drugs overlap with the hotspots for some of the disease-causing FAD mutations (Fig. 4, https://www.alzforum.org/mutations) *(36, 37)*. The overlapping structural elements are in line with the activity studies which showed that the drugs and FAD mutations can produce similar changes in γ-secretase activity *(6, 10, 12)*.

The possibilities that several drugs can bind to γ-secretase simultaneously have been indicated in some of the earlier studies *(43)*. Multiple studies also indicated that the same drugs can sometimes bind to different sites *(37, 42-45)*. However, to our knowledge, the functional consequences of multiple enzyme-drug interactions have been explored only in our mechanistic studies of biphasic dose-response *(10, 11)*.

### All-atom molecular dynamics studies of binding interactions between biphasic-drugs and γ-secretase in the presence and absence of substrate (Fig. 5-6)

We have combined molecular docking studies (Fig. 4) with the molecular dynamics studies (Fig. 3), with a desire to describe how drugs can affect the dynamic changes in presenilin structure that regulate enzyme-substrate interactions (Fig. 3) *(36, 37)*. We found that all four drugs can decrease the mobility of the flexible protein loops that regulate the opening of the active site tunnel, or the release of tripeptide catalytic intermediates in processive catalysis (Figs. 5-6) *(6, 12, 25)*. DAPT has the smallest effect on the decrease in mobility as illustrated by RMSF values, whereas avagacestat and LY-411,575 have the biggest effect (Figs. 5-6 panel D). Thus, the changes in mobility correlate with the measured binding affinities (Fig. 1).

**Figure 5.**
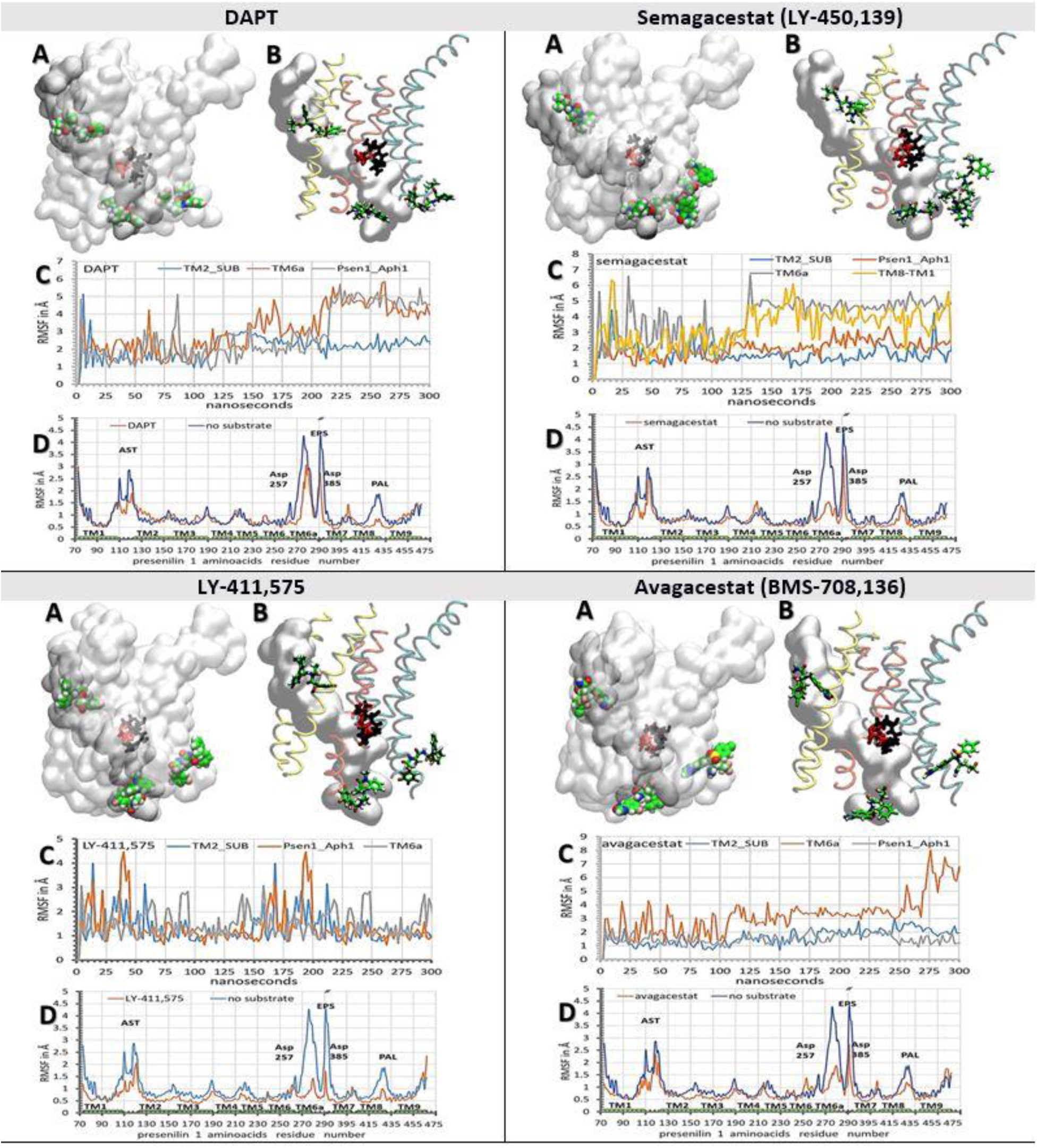
(A-D). Binding interactions between the biphasic-drugs and γ-secretase in complex with its substrate (PDB:6IYC) *(36)*. We used all-atom molecular dynamic calculations *(65)* to describe how different biphasic-drugs can affect dynamic interactions between γ-secretase and its substrate *(36, 37)*. The molecular dynamic calculations *(65)* started with γ-secretase structures in complex with biphasic-drugs that have been prepared in molecular docking studies (Fig 4) *(46)*. The results are presented using models of presenilin structure (A-B) *(75)* and quantitatively using RMSF values for drugs (C) and for presenilin (D) *(38, 39)*. (**<u>A</u>**) The presenilin structures are shown as transparent surfaces to make structures underneath the surface visible. (**<u>B</u>**) TM2 and TM3 are shown in yellow, TM6, TM6a, and TM7 are shown in orange, and TM8, TM9, and TM1 are shown in cyan. (**<u>A</u>**-**<u>B</u>**) Buried under the surface in the active site tunnel is the substrates (gray surfaces), the active site Asp 257 and Asp 385 (red licorice), and PAL motif (black licorice, Pro 435, Ala 434, Leu 433). Drug molecules are shown as: carbon green, oxygen red, nitrogen blue, fluorine, and chlorine as pink. (**<u>C</u>**) RMSF values for drugs bound at different sites are shown as a function of molecular time *(65)*. The steep increases in RMSF values indicate the sliding of the drugs in the binding sites, while the fluctuations in RMSF values indicate the relative mobility of the drugs in their binding sites. (**<u>D</u>**) RMSF values for individual amino acids were used to show how drugs can decrease protein mobility at different sites *(38, 39)*. Different structural elements are mapped on the graph: TM1-TM9 <u>T</u>rans-<u>M</u>embrane helix 1 to 9, AST membrane-embedded opening of the <u>A</u>ctive <u>S</u>ite <u>T</u>unnel, EPS <u>E</u>ndo-<u>P</u>roteolytic <u>S</u>ite, PAL motif (Pro 435, Ala 434, Leu 433) *(40)*, and Asp 257 and Asp 385 in the active site.

The most significant observation is that the drugs can cooperate in penetration into the active site tunnel in the absence of the substrate (Fig. 6, supp. Fig. 3, supp. video 1). γ-Secretase structures with the drugs bound simultaneously at each end of the tunnel have been compared with the structures that had the drugs bound only at one end (supp. Fig. 3). We found that the drugs penetrated faster and deeper in the active site tunnel when they bind simultaneously at each end of the tunnel (Fig. 6, supp. Fig. 3, supp. video 1). These results are in agreement with some of the earlier studies which indicated that the binding of one drug molecule can facilitate the binding of another drug molecule *(43)*. DAPT and LY-411,575 showed the deepest penetration (Fig. 6, supp. Fig. 3, supp. video 1). The two drugs can reach the active site Asp 257 and Asp 384 from both ends of the tunnel (Fig. 6, supp. Fig. 3, supp. video 1). At the cytosolic end the drugs can penetrate in the active site tunnel driven by the highly dynamic endoproteolytic site between TM6, TM6a, TM7. The penetration in the active site tunnel between TM2 and TM3 is driven by a sequence of specific interactions. Buried in a predominantly hydrophobic cavity, the drugs form at first several hydrophobic interactions, most notably with Ile 138 and Val 142 on TM2. Phe 237 and Tyr 115 can form π-π stacking interactions with aromatic parts of each drug (supp. Fig. 2). The inhibitors can also form dynamic hydrogen bonds with the side chains on Thr 44 on the substrate and with Tyr 115 on presenilin (supp. Fig. 2). The site is dominated by hydrophobic interactions. In less than 10% of molecular dynamic conformations, the drugs form one hydrogen bond with the protein. DAPT and LY-411,575 can actively penetrate in the active site tunnel driven by their aromatic rings that can form π-π stacking interactions with Phe 237 and Tyr 115 in the tunnel (Fig. 2, supp. Fig. 2). The penetration is much slower with semagacestat and avagacestat, because these drugs lack the flexible aromatic structure that can support optimal interactions.

**Figure 6.**
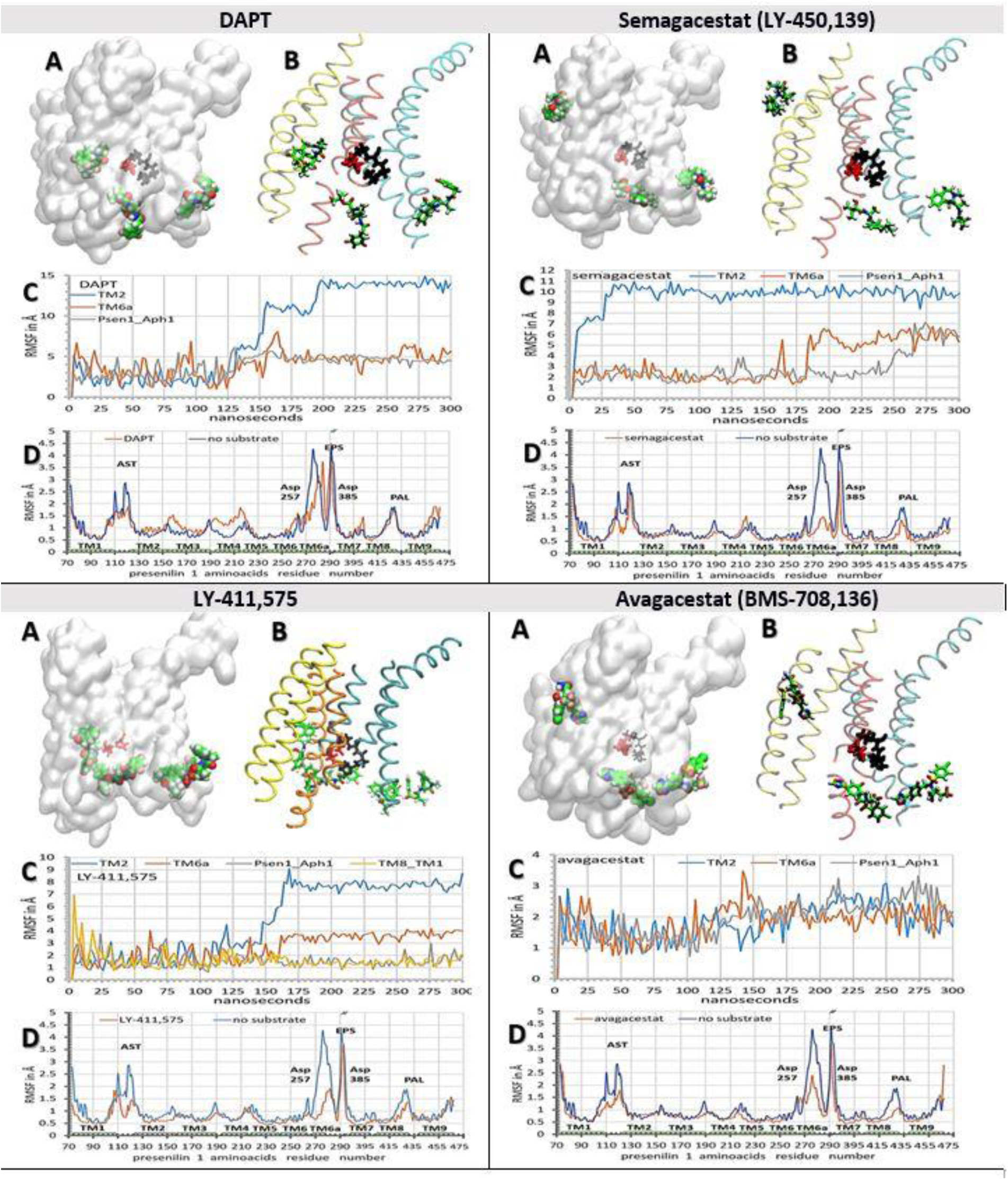
(A-D). Binding interactions between the biphasic-drugs and γ-secretase in the absence of substrate. We used all-atom molecular dynamics calculations to analyze whether molecules of biphasic-drugs can penetrate in the active site tunnel in the absence of the substrate *(36, 37)*. The results are described using surface (A) and ribbon (B) models of presenilin 1 *(75)*, and quantitatively using RMSF values for the drugs (C) and presenilin (D) *(38, 39)*. The molecular dynamics calculations *(65)* started with γ-secretase structures in complex with different biphasic-drugs that have been calculated in molecular docking (Fig 4) *(46)*. **(<u>A</u>)** The presenilin structures are shown as transparent surfaces to shown the structures below the surface. **(<u>B</u>)**TM2 and TM3 are shown in yellow, TM6, TM6a, and TM7 are shown in orange, and TM8, TM9, and TM1 are shown in cyan. (**<u>A</u>**-**<u>B</u>**) The figures show how different drugs can penetrate in the active site tunnel to a different extent in the absence of the substrate. Buried under presenilin surface are active site Asp 257 and Asp 384 (red licorice) and PAL motif (Pro 435, Ala 434, Leu 433, black licorice) *(40)*. Drug molecules are shown as carbon green, oxygen red, nitrogen blue, fluorine, and chlorine as pink. (**<u>C</u>**) RMSF values for drugs bound at different sites are shown as a function of molecular time *(65)*. The steep increases in RMSF values indicate the moments when the drugs are sliding in their binding sites, while the fluctuations in RMSF values indicate the mobility of the drugs in their binding sites. (**<u>D</u>**) Average RMSF values for individual amino acids were used to illustrate changes in protein mobility caused by the drugs *(38, 39)*. All four drugs can decrease mobility at the specific sites in the presenilin structure to a different extent. Different structural elements are mapped on the graph. TM1-TM9 Trans-Membrane helix 1-9, AST membrane-embedded opening of the <u>A</u>ctive <u>S</u>ite <u>T</u>unnel, EPS <u>E</u>ndo-<u>P</u>roteolytic <u>S</u>ite, PAL motif (a.a. 433 to 435) *(40)*, and Asp 257 and Asp 385 in the active site.

γ-Secretase structures with drug molecules bound only at the cytosolic end of presenilin structure showed that drugs can bind and slide between dynamic protein loops that form 18 Å long surface from TM6 to TM8 (Figs. 5-6). The drugs can bind between TM6 and TM7 and slide toward the active site Asp 257 and Asp 385, or bind between TM7 and TM8 and slide towards the PAL motif (a.a. 433-436, *(40)*). The drugs that bind and slide between TM8 on presenilin and the Aph1 subunit can stabilize the gap between the two proteins in the open position (Figs. 5-6). The most extensive sliding is observed with DAPT and semagacestat, the lowest sliding is observed with LY-411,575 and avagacestat (Figs. 5-6, supp. Fig. 3, supp. video 1). The sliding drugs form mostly hydrophobic interactions with the dynamic cavities on the presenilin surface and up to 3 highly dynamic hydrogen bonds with the flexible protein loops or the surrounding water molecules. In the absence of the substrate, the flexible protein loops drive the drug in the position of the substrate between TM6, TM6a, and TM7 (Fig. 6). In the presence of the substrate, the substrate is forcing the drugs away from the tunnel towards TM8 and Aph1 (Fig. 5). Binding of the first drug molecule can facilitate the binding of the second drug molecule by opening the protein loops at the adjacent sites. Docking studies showed that with all four drugs as much as 3 molecules can bind into the 18 Å long surface in the absence of the substrate (Fig. 6). Only LY-411,575 can form stable interactions with 3 molecules bound, while with the other drugs only two molecules can bind in parallel at this end. These results are consistent with Hill’s coefficients in the biphasic dose-response curves (Fig. 1). The binding of multiple drug molecules can block drifting within the drug-binding cavities, and thus multiple drug molecules compete more effectively with the conformational changes in the flexible protein loops (Fig. 6).

The RMSD values for all four drugs showed the most stable binding in the hydrophobic gap between the Aph1 subunit and TM8 in presenilin (Figs. 5-6). The substrate has the smallest effect on this site (Figs. 5-6). The drugs facilitate the breaking of the contacts between TM8 on presenilin and TM2 and TM3 on Aph1 (Figs. 5-6). The drugs are buried in a hydrophobic gap between Leu 420, Leu 423, Ile 427 on TM8 on presenilin and Leu 93, Leu 96 on TM2 of Aph1. The dynamic conformational changes can drive the gap between open and closed conformation. The closed conformation is stabilized by a salt bridge between Glu367 at the endoproteolytic site and Lys 429-430 at the cytosolic end of TM8.

With all four drugs, γ-secretase can bind one drug molecule at the membrane end of the active site tunnel *(37)* (Figs. 5-6). In the absence of the substrate, the drugs form contacts with TM2, TM3, and TM5 (Fig. 6 A-B). The mobile linkers between TM2, TM3, and TM5 (Fig. 3, residues 110 to 130, and 214 to 217) control the width of the cavity and the penetration of the drugs in the active site tunnel (Fig. 6, supp. Fig. 3). The drugs form hydrophobic interactions with Ile 138 and Val 142 on TM2, and π-π stacking interactions between the aromatic rings and Phe 237, and Tyr 115 (supp. Fig. 2). The substrate can displace the interactions between the drugs and TM3 and TM5 (Fig. 5 A-B, supp. Fig. 2). The drug is burred in a tight hole between TM2 and Gly 38 and Val 40 on the substrate (Fig. 5, supp. Fig. 2). This observation is in agreement with the studies that indicated that the drugs can form nonspecific interactions with substrate and affect Aβ production *(47)*.

With all four drugs, the binding constants in cell-based assays (Fig. 1) correlate with the observed RMSF values (Figs. 5-6 panel C). The conformational mobility within the binding sites appears to be the main determinant of the binding affinity (supp. video 1). DAPT has the weakest binding affinity (Fig. 1) and the highest RMSF values (Figs. 5-6). DAPT has an elongated flexible structure that can effectively slide in the cavities on the protein surface (Fig. 3, supp. Fig. 3, supp. video 1). DAPT and LY-411,575 showed a very similar binding mechanism and the two drugs show the deepest penetration in presenilin structure (Fig. 5-6 and supp. Fig. 3). LY-411,575 like DAPT can extend in the cavities on the presenilin surface. However, LY-411,575 has one bulky end that can anchor the molecule at one site and that can make it more resistant to the displacement by the conformational changes in the presenilin structure (Supp. Figs. 2-3). Semagacestat has some of the same functional groups as DAPT and LY-411,575 (Fig. 2), however as the smallest molecule (Fig. 2), it has much less influence on the conformational changes in the presenilin structure (Fig. 5-6, supp. Fig. 3, supp. video 1). For those reasons, semagacestat cannot penetrate in the active site tunnel as much as the bigger LY-411,575 and DAPT molecules (supp. Fig. 3, supp. video 1). Small semagacestat molecules (Fig. 2) can be readily displaced by the substrate at each end of the active site tunnel (Fig. 5 panel A, and Fig. 6). Avagacestat can resist the flexible changes in the protein structure because it has the widest structure with several rigid spots (Fig. 2), which allow the molecule to extend and resist to the conformational changes in presenilin structure (Figs. 5-6, supp. Fig. 3, supp. video 1). Avagacestat cannot slide on the protein surface and shows on average the lowest RMSF values (Figs. 5-6 panels C-D). LY-411,575 and avagacestat depend on different mechanisms to resist the forces caused by the conformational changes in the flexible protein parts. The two molecules show the lowest RMSF values and the strongest binding constants (Fig. 1 and Figs. 5-6).

### All-atom molecular dynamic studies of γ-secretase structure with Aβ catalytic intermediates in processive catalysis

Structures of γ-secretase in complex with different Aβ catalytic intermediates were used for the analysis of modulation of Aβ production by biphasic-drugs *(6, 8, 12, 25)* (Fig. 7 and supp. Fig. 1B). Earlier studies of biphasic dose-response with DAPT showed that the drugs can facilitate the production of shorter Aβ products at the lower concentrations, and production of longer Aβ peptides at higher concentrations *(12)*. Those observations indicated that with different Aβ catalytic intermediates the drugs bind at different sites. A higher affinity site with the shorter Aβ products and lower affinity sites with the longer Aβ products.

**Figure 7.**
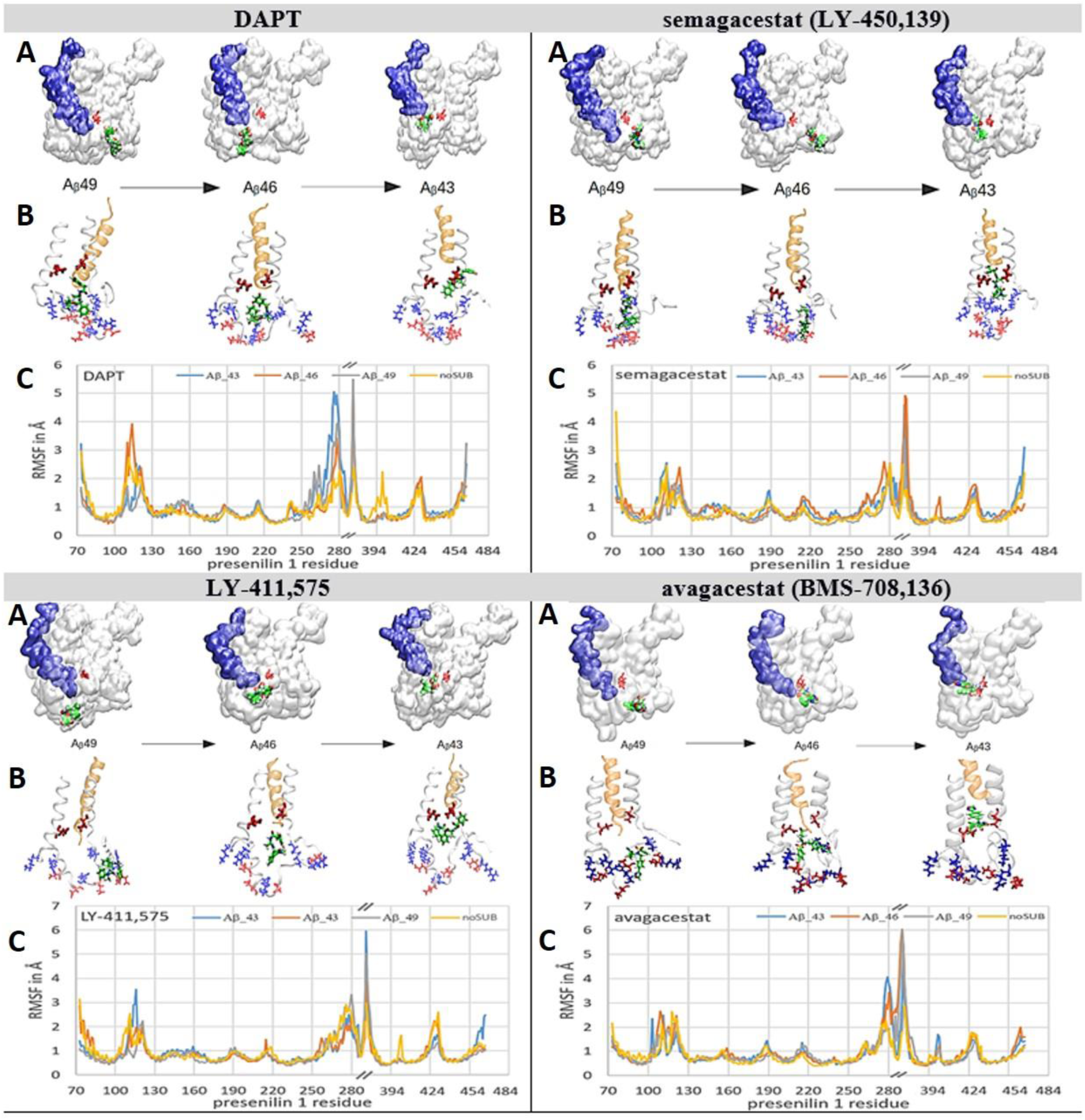
(A-C). Biphasic-drugs can bind to γ-secretase and selectively interfere with the processive proteolytic cleavages. We used all-atom molecular dynamics studies *(65)* to analyze whether molecules of biphasic-drugs can penetrate into the active site tunnel when γ-secretase is in complex with different Aβ catalytic intermediates. With all four drugs, the deepest penetration is observed with Aβ 43, the lowest penetration is observed with Aβ 49. The penetrations are depicted using the presenilin structures *(36)*, and quantitatively using RMSF values as a function of amino acid positions *(38, 39)*. (**<u>A</u>**) The presenilin structures are shown as a white transparent surface to make the structures below the surface visible. Buried underneath the surface in the active site tunnel are different Aβ catalytic intermediates (blue surface) and active site Asp 257 and Asp 385 (red licorice). The drugs are shown as green VanderWaals models. (**<u>B</u>**) The white ribbon models depict amino acids 240 to 394 in presenilin structures. The gold ribbon models depict different Aβ catalytic intermediates. The positively charged amino acids are shown as blue licorice, negatively charged amino acids are shown as red licorice, including the active site Asp 257 and Asp 385. Drugs are depicted as green licorice models. (**<u>C</u>**) RMSF values as a function of amino acid positions show how biphasic drugs can affect different structural parts to a different extent with different Aβ catalytic intermediates *(38, 39)*.

We have prepared γ-secretase in complex with Aβ 40, Aβ 43, Aβ 46, and Aβ 49, for analysis of the binding interactions with all four biphasic-drugs (Fig. 7). The structures with Aβ 40, Aβ 43, Aβ 46, and Aβ 49 represent to different extent a mixture between *open* and *closed* structures of γ-secretase *(36, 37)*. When Aβ substrate of different length are bound in the active site tunnel, the tunnel is predominately closed at its cytosolic end (Fig. 7, supp. Fig. 1B). Thus, we started the simulations by docking the drugs at the cytosolic end of presenilin with the active site tunnel closed.

The drugs penetrate into the active site tunnel in the presence of Aβ catalytic intermediates (Fig. 7) driven by the same mechanism that drives the penetration of multiple drug molecules at different ends of the tunnel (Fig. 6, supp. video 1, sup Fig. 3). Aβ molecules can facilitate opening of the tunnel and penetration of the drugs into the tunnel (Fig. 7) faster than the drug molecules that bind at the membrane-embedded end of the tunnel (Fig. 6, supp. video 1, sup Fig. 3). The drugs that penetrate into the active site tunnel bind between flexible loops that can selectively disrupt processive cleavages of the nascent Aβ peptides (supp. Fig. 1, *(6, 8, 12)*). Different binding interactions, and different RMSF values, are in agreement with the earlier observations that modulation of Aβ production by different drugs depends on the structure and concentration of each drug *(6, 8, 12, 25)*.

With short Aβ 40 and Aβ 43, the drugs can penetrate deep into the active site tunnel and show the lowest RMSF values when bound inside the tunnel (Fig. 7). The drugs bind between amino acids 146 to 173 and 226 to 387 in the predominantly hydrophobic surface that can form only up to one dynamic hydrogen bond (Fig. 7). Parts of the binding surfaces are Pro284, Ala285, and Leu286, and aromatic amino acidsTyr154, Trp165, and Phe283 (Fig. 7). Aromatic residues can contribute to π-π stacking. The penetration depends on the molecular structures. DAPT and LY-411,575 have an elongated flexible structure that can line up with the tunnel walls and penetrate deeper in the channel. Lower depth of penetration is observed with semagacestat. Semagacestat has structural similarities with DAPT and LY-411,575, but its small structure has much smaller effects on flexible loops in presenilin structure. Avagacestat has a wide structure that cannot line with the tunnel walls and penetrate deep in the tunnel.

The longer Aβ 46 and Aβ 49 catalytic intermediates do not allow deep penetration of the drugs in the active tunnel (Fig. 7). The drugs bind in the hydrophobic cavities where they can form up to 3 dynamic hydrogen bonds with the charged amino acids on dynamic flexible loops (Fig. 7, sup. Fig. 1). The drugs bind between amino acids 73 to 83, 378 to 381, and 417 to 435 in the active site loop. Part of the binding site is PAL motif (Pro433, Ala434, Leu435, Pro436) *(40)*, and aromatic amino acids Tyr77 and Phe428 that can result in π-π stacking interactions with the drug (Fig. 7).

## Discussion and Conclusions

### Structural insights in the molecular mechanism behind biphasic dose-response curves

We used molecular structures to extend our earlier mechanistic studies of biphasic dose-response in measurements of γ-secretase activity *(7, 10, 11)*. The presented structural studies are in agreement with conclusions from the earlier activity studies with biphasic-drugs *(6, 8, 10-12, 15)*. Multiple drug molecules that bind to γ-secretase can produce the biphasic dose-response by selectively acting on different steps in the catalytic cycle (Figs. 3-7); from the initial substrate binding (Fig. 6, supp. Fig. 3, supp. video 1), to the final processing and the release of Aβ catalytic intermediates (Figs. 5 and 7).

Combined structure-activity studies showed that the biphasic-drugs can activate γ-secretase by forcing the active site tunnel to open (supp. Fig. 3, supp. video 1), when the rate-limiting step is the tunnel opening *(10, 37)*, and formation of the enzyme-substrate complex *(10, 36)*. The enzyme activity studies showed that the drugs can activate γ-secretase by facilitating enzyme-substrate interaction at limiting substrate conditions *(10)*. The structural studies showed that the active site tunnel is tightly closed in the absence of the substrate to prevent water and ions from leaking across the membrane *(36, 37)* (Fig. 3, supp. Fig. 1B). The opening of the tunnel is the rate-limiting step in the initial enzyme-substrate recognition *(36, 37)* (Fig. 3). Thus, the drugs that can force the opening of the tunnel can activate γ-secretase when the tunnel opening and enzyme-substrate recognition control the rate limiting-step. The active site tunnel is predominantly closed in the absence of substrate due to hydrophobic interactions inside the tunnel and due to contraction between flexible protein loops that control the ends of TM domains (Figs. 3, supp. Fig. 1-3) *(36, 37)*. The drug-molecules can bind to the loops and force the TM domains apart by mimicking the substrate in the process of enzyme-substrate recognition (Fig. 6, Supp. Figs. 2-3). The highest activation is observed with LY-411,575, which has the strongest binding affinity and thus the best chance to stabilize the open tunnel structure (Fig. 1) *(34)*. LY-411,575 has an elongated flexible structure (Fig. 2) that can penetrate deep in the tunnel (Fig. 6 and supp. Fig. 3) where its bulky head can form π-π stacking interactions (supp. Fig. 2) and stabilize the presenilin structure in its open conformations *(36, 37)*. The active site tunnel and the flexible protein loops on the cytosolic side of the presenilin structure (Fig. 3), resemble the functions of the active site loops that are usually targeted in drug-development studies with soluble proteases *(48, 49)*.

Combined structure-activity studies showed how biphasic-drugs can inhibit γ-secretase. The structural studies showed that the drugs bind to different sites on γ-secretase in the presence and the absence of substrate (Fig. 5-7). In the absence of the substrate, the drugs bind in place of the substrate and to a different extent penetrate into the tunnel (Fig. 6, supp. Fig. 3, supp. video 1) *(36, 37)*. Such binding can explain why studies of γ-secretase activity showed competition between the drugs and the substrate in activation of γ-secretase *(8, 10)*. The structural studies further showed that in the presence of the substrate, the drugs bind to the flexible protein parts that participate in the processive catalysis (Fig. 3 and Fig. 5). The substrate cannot displace the drugs from those sites (Fig. 5). These observations are in agreement with the activity studies which showed that the biphasic-drugs are uncompetitive inhibitors *(10, 11)*. The uncompetitive inhibitors are not acceptable in any of the drug-development strategies with γ-secretase as the target. At lower doses, the uncompetitive inhibitors can produce the same changes in γ-secretase activity as FAD mutations in presenilin 1 and possibly facilitate pathogenesis *(7, 10, 24)*. At higher doses, the uncompetitive inhibitors can stop the vital functions of γ-secretase in cell physiology *(2, 4, 50)*. During the catalysis, the flexible protein loops have to drive processive cleavages and the release of tripeptide intermediates *(12, 25)* (Fig. 3). The regulation of processive catalytic steps can be a crucial mechanism in pathogenesis *(12, 21, 24, 25)*.

Combined structure-activity studies showed that the biphasic-drugs can modulate the production of Aβ catalytic intermediates by penetration into the active site tunnel to different depth with different binding affinity. The activity studies showed that different Aβ products showed biphasic dose-response at different drug concentrations *(12, 25)*. The shorter Aβ products show activation at the lower drug concentrations while the longer Aβ products show activation at the higher drug concentrations *(12, 25)*. Thus, there are high-affinity sites for the drugs with the shorter Aβ peptides, and the lower affinity sites for the drugs with the longer Aβ peptides. In line with those observations, we found that the drugs can penetrate deeper into the presenilin structure with the shorter Aβ peptides (Fig. 7). The drugs can mimic tripeptide catalytic products in the processive cleavages (Figs. 3, Fig. 7, supp. Fig. 1). The depth of penetration, and consequently the binding affinity, depends on the ability of each drug to jam the flexible loops that regulate processive catalysis and the tripeptide release (Fig. 7). It is very likely that all studies that have reported modulation of Aβ production, or selectivity between APP vs. Notch substrate, had the biphasic response mechanism that was not detected because the measurements used limited concentration range for the drugs and substrate *(8, 10, 12, 25)*.

The drugs and the FAD mutations target the same dynamic processes in presenilin structures (Figs. 4-7, and https://www.alzforum.org/mutations) *(3, 36, 37)*. FAD mutations can affect the biphasic dose-responses *(7)*. Selective action on different steps in the catalytic cycle (Fig. 3-7) can explain how biphasic-drugs and different FAD mutations can both activate and inhibit γ-secretase *(6-8, 10, 12, 15, 21, 24, 51)*. The selective interference with the dynamic conformational changes at the cytosolic end of TM6, TM6a, TM7 (Fig. 7, supp. Fig. 1) can explain how biphasic-drugs and FAD mutations can favor increase in the production of the longer more hydrophobic Aβ catalytic intermediates *(6, 8, 10, 12, 15, 21, 24, 25)*.

### Future drug development strategies with γ-secretase as the target enzyme

The biphasic dose-response curves can give a false impression that the drugs can be used at precise doses only as activators or only as inhibitors of γ-secretase (Fig. 1) *(9, 14, 15)*. Such targeted dose therapy is not possible, because, both the activation and the inhibition by biphasic-drugs can decrease the catalytic capacity of γ-secretase just like the FAD mutations *(7, 10)*. The decrease in catalytic capacity can be a result of the increase in enzyme saturation with its substrate, or a decrease in the turnover rates, or both *(7, 10, 16)*. Very diverse studies showed a good correlation between a decrease in γ-secretase capacity to process its substrates and the pathogenic events *(7, 26, 51-59)*. Based on those observations we are proposing that competitive inhibitors of γ-secretase have the best chance to become drugs for Alzheimer’s disease *(7, 16, 18)*. Similar to the competitive inhibitors, the protective A673T mutation in the APP substrate can decrease the extent of enzyme saturation with its amyloid substrate with no effects on the turnover rate constant *(51)*. The aim is not to inhibit γ-secretase. The aim is to modulate the extent of the enzyme saturation with its substrates in correlation with changes in the metabolic load for APP and Notch substrates *(7, 16)*.

The development of competitive inhibitors requires changes in the drug-screening strategies *(16, 18)*. The majority of the past drug development studies did not pay attention to the extent of γ-secretase saturation with its substrate, and most often, the measurements used saturating substrate. Such assays are the cheapest and the fastest. However, such assays can detect only uncompetitive inhibitors that can produce the same changes in the γ-secretase activity as FAD mutations in presenilin 1 *(7, 10, 21, 24)*. In cells, in physiological conditions, γ-secretase is far from saturation with its substrate *(7-9)*. The sub-saturated enzymes are a fundamental physiological mechanism for all enzymes *(20)* that can assure the fastest and best-controlled response to metabolic fluctuations *(16, 18, 19)*. The artificial preclinical studies with γ-secretase at saturating substrate have limited clinical relevance and often lead to misleading conclusions *(6, 8, 9, 15)*. Measurements with poorly defined saturation of γ-secretase with its substrate can explain why so many studies reported inconsistent and irreproducible results on modulation of Aβ production, and/or on selectivity between different substrates *(4)*. Different substrates and different Aβ products have different Kӎ values *(12, 16, 21, 22)*. Kӎ values are crucial parameter for a meaningful description of preferences for different substrates and Aβ products *(16, 18, 21, 22)*.

Future drug-development strategies can also target the Aph1 subunit (Fig. 4). Different Aph1 subunits can affect the production of toxic Aβ products, with no effect on turnover rates of γ-secretase *(60)*. The mechanism is still unknown *(60)*. We found that all four drugs target the gap between presenilin and Aph1 subunits (Figs. 5-6). The function of the gap is unknown, some studies have suggested that the gap could be a water channel for the active site *(36)*.

### Closing remarks

The initial “fast-and-cheap” drug-screening strategies with γ-secretase as the target enzyme must be supplemented with the fundamental protocols for analysis of enzyme activity *(16-18)* to achieve reproducible results and sustained progress *(10, 12, 16, 18, 22, 25, 26)*. The expensive failures in clinical trials with semagacestat *(6, 15, 29, 30, 61)* have been much smaller with avagacestat *(9, 27)*, because, the preclinical studies with avagacestat were much better documented and comprehensive in design *(8, 9, 14, 27, 28)*.

We further point out that additional unique opportunity with γ-secretase is that the mechanistic studies can be used for development of compounds that can produce the same type of pathogenic changes in γ-secretase activity as disease-causing FAD mutations *(6, 7, 10, 12, 21, 23, 24, 26, 51-58)*. Such compounds can be a powerful tool for the development of exceptionally precise protocols for analysis of different causes of pathogenic changes in γ-secretase activity in cell cultures and model animals *(23)*.

The future structural studies of γ-secretase can use tight-binding drugs such as LY-411,575 to capture the structure of dynamic conformational changes that regulate substrate binding and catalysis *(36)* (Fig. 6, supp. Fig. 3, supp. video 1).

## Materials and Methods

### Materials

The drugs were purchased from Calbiochem: <u>DAPT</u> (*N*-[*N*-(3,5-difluorophenacetyl)-L-alanyl]-*S*-phenylglycine *t*-butyl ester). <u>Semagacestat</u>, LY-450,139 2S)-2-Hydroxy-3-methyl-N-[(1S)-1-methyl-2-oxo-2-[[(1S)-2,3,4,5-tetrahydro-3-methyl-2-oxo-1H-3-benzazepin-1-yl]amino]ethyl]-butanamide; <u>LY-411,575</u>, N2-[(2S)-2-(3,5-Difluorophenyl)-2-hydroxyethanoyl]-N1-[(7S)-5-methyl-6-oxo-6,7-dihydro-5H-dibenzo[b,d]azepin-7-yl]-L-alaninamide; Avagacestat, (2R)-2-[N-[(4-Chlorophenyl)sulfonyl]-N-[[2-fluoro-4-(1,2,4-oxadiazol-3-yl)phenyl]methyl]amino]-5,5,5-trifluoropentanamide, (R)-2-(4-Chloro-N-(2-fluoro-4-(1,2,4-oxadiazol-3-yl)benzyl)phenylsulfonamido)-5,5,5-trifluoropentanamide, BMS-708163.

### Secretion of Aβ 1-40 in cultures of SHSY5 cells in the presence of increasing concentrations of drugs

SHSY5 cells were purchased from ATCC as passage 11 and maintained in Dulbecco’s modified Eagle’s medium (DMEM) supplemented with 10% fetal bovine serum. The measurements of the biphasic response in the presence of drugs and the corresponding data analysis have been described in detail in our earlier studies *(7, 10)*. Briefly, different concentrations of drugs were prepared in DMSO, and added to the cells so that the final DMSO concentration in the culture was 0.1% (v/v). DMSO vehicle represents 0 nM drugs. The cells were incubated in 6 well-plates, with the drugs at given concentrations between 12-18 hours. We took maximal attention to measure all four drugs under identical conditions. The same batch of cells was used in parallel for measurements with all four drugs. Identical conditions were used for the incubation of drugs with the cells, sample harvesting, and measurements of Aβ 1-40.

### Sandwich ELISA for quantitative detection of Aβ 1-40

The assays closely followed the manufacturer’s instructions. Sandwich ELISA kits for quantitative detection of human Aβ 1-40 peptides in a flexible 96 well format were purchased from Millipore (cat. #. TK40HS, The Genetics company Switzerland). The assay had a linear response in the range from 6 - 125 pM of Aβ 1-40. Aβ 1-40 samples from cell cultures were used immediately after collection following the manufacturer suggestion and our earlier reported experimental experiences *(10)*. The wells were filled with 50 µl of the antibody conjugate solution and 50 µl of the sample. The Aβ 1-40 standards supplied by the manufacturer were prepared in parallel with the other samples. The prepared wells were wrapped in aluminum foil and incubated overnight at 4 °C with gentle mixing. The next day each well was washed five times with 300 µl of wash solution. After each 20 minutes wash, the wash solution was poured out and the wells were dried by tapping the plates on an absorbing paper. After washing the wells were filled with 100 µl of the enzyme conjugate solution, covered, and incubated for 30 min at room temperature with shaking. The washing procedure was repeated once again to remove the excess of the enzyme-conjugate. Next 100 µl of the substrate solution was added in each well in dark and kept for 30 minutes covered at room temperature. The reaction was quenched by adding 50 µl of stop solution to each well, and within 15 min the signal intensity was read by measuring absorption at 450 nm.

### Inhibitor docking studies

Binding sites for semagacestat, avagacestat, LY-411,575, and DAPT were calculated using RxDock and AutoDock Vina 1.1.2. following the standard protocols *(46, 62)*. Briefly, the ligands were hydrogenated and charged using the Gasteiger protocol and pH=7.0. Proteins were protonated at pH=7.0 using AMBER98S force filed with Asp 257 and Asp 385 unprotonated.

### Residue based Coarse-Grained Molecular Dynamics

Coarse-grained molecular dynamics calculations with γ-secretase structure within the lipid bilayer were prepared using CHARMM-GUI 32 Martini Bilayer Maker 33 *(35, 63)*. OPM protocol was used to position and orients proteins in a lipid membrane bilayer *(64)*. The systems were relaxed using equilibration steps with the temperature set to 310 K using V-rescale coupling, and the pressure was set to 1.0 bar using semi-isotropic Berendsen coupling *(65)*. The systems with mixed lipid bilayer have 1355 residues, 1604 lipid molecules, 71112 water molecules, 924 Na+ ions, 791 Cl-ions in a 210 Å x 210 Å x 246 Å box. Lipid composition of membrane bilayer in CG systems *(66)* (lipid type, number of lipid molecules, percentage of lipid molecules): phosphatidylcholine (POPC), 340, 21%; phosphatidylethanolamine (POPE), 176, 11%; phosphatidic acid (POPA), 16, 1%; phosphatidylserine (POPS), 64, 4%; sphingomyelin (PSM), 96, 6%; phosphatidylinositol (POPI), 32, 2%; cholesterol (CHOL), 880, 55%. The lipid composition is crucial for achieving presenilin structure with proper orientation between the active site aspartates *(67)* and (Svedružić *et. al*. in preparation). γ-Secretase structures in molecular dynamics simulations are depicted in time-steps of 20 fs to describe periods of 10 to 20 µs of molecular events in 1000 to 2000 frames.

### All-atom molecular dynamics studies

γ-Secretase structures in complex with different drugs from molecular docking studies *(46, 68)* were used as the starting structures for molecular dynamic studies. γ-Secretase structures with the active site tunnel closed were prepared in molecular docking studies using the structures without the substrate as the starting structures.

All-atom simulations of γ-secretase in lipid bilayer were prepared using CHARMM-GUI Membrane Builder 35–37 *(69)*. OPM protocols were used to position and orients proteins in a lipid membrane bilayer *(64)*. The systems were relaxed using a sequence of equilibration steps with the temperature set to 303.15 K using Nose-Hoover coupling, and the pressure was set to 1.0 bar using semi-isotropic Parinello-Rahman coupling. The systems with mixed lipid bilayer have 1309 residues, 708 lipid molecules, 132325 TIP3 water molecules, 422 Na+ ions, and 369 Cl-ions in a 144 Å x 144 Å x 247 Å box. Lipid composition of membrane bilayer in AA systems with mixed lipid bilayer (lipid type, number of lipid molecules, percentage of lipid molecules): phosphatidylcholine (POPC), 152, 21%; phosphatidylethanolamine (POPE), 78, 11%; phosphatidic acid (POPA), 8, 1%; phosphatidylserine (POPS), 28, 4%; sphingomyelin (PSM), 42, 6%; phosphatidylinositol (POPI), 14, 2%; cholesterol (CHOL), 386, 55%.

The systems used in our simulations contain 85080 TIP3 water molecules, 272 POT ions, 221 CLA ions, and 708 lipid molecules. The overall number of atoms in these systems is 347516 and they are contained in a 144 Å x 144 Å x 176 Å box. The temperature of the simulation was set to 303.15 K using Nosé–Hoover coupling. The pressure was set to 1 bar using semi-isotropic Berendsen coupling. The constraint algorithm was LINCS and the cut-off scheme was Verlet. To ensure proper system relaxation, one minimization step and six equilibration steps were used. The simulations analyzed molecular processes from 300 to 600 nanoseconds on a molecular time scale with 2 fs time step.

The validities of presenilin structures in each of the molecular dynamic simulations were analyzed by comparing the active site structures with the mechanistic studies of the active site of aspartic protease*(67)* and by comparing the calculated and experimental pKa for the active site aspartates *(70, 71)* (Svedružić et. al. manuscript in preparation).

The ligand parameterization was prepared using ACPYPE tools *(72)* or the CHARMM-GUI Ligand Reader & Modeler, to calculate CHARMM-compatible topology and parameter files *(73)*. All-atom input files with γ-secretase in lipid bilayer were prepared using the CHARMM-GUI Membrane Builder *(74)*. EM refined structures of γ-secretase (PDB: 6IYC) *(36)* had a total of 1355 residues, 5 chains, and a resolution of 2.60 Å. The proteins were positioned and oriented in lipid membrane bilayers using OMP protocol*(64)*. The molecular structures for semagacestat, avagacestat, LY-411,575, and DAPT were taken from the ChemSpider and PubChem databases. All simulations used GROMACS version 2019.4 *(65)*. Molecular dynamics for γ-secretase in complex with different drugs were depicted in time-steps of 2 fs to represent a total of 300 to 600 nanoseconds of molecular events in 100 to 200 recorded frames.

### Data analysis and presentation

Molecular imaging and analyses molecular properties using VMD 1.9.3 and UCSF Chimera 1.14 *(68, 75)*. The molecular trajectories have been analyzed using Bio3D package with R 3.6.2 and Rstudio 1.2.5019’s *(38, 39)*. RMSF (root-mean-square-fluctuations) values were calculated as a function of molecular time*(65)* or the position of amino acid residues *(38, 39)*. All biphasic profiles were analyzed using nonlinear regression and the equation for the biphasic dose-response curve that was described in detail in our earlier studies *(7, 10)*:

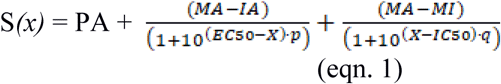

where, *S(x)* represents measured activity at inhibitor concentration *x. PA* is the physiological Aβ 1-40 production activity at inhibitor concentration zero. *MA* is the calculated maximal activity caused by activation if there is no competing inhibition. *MA* values roughly correlate with the capacity of drugs to induce increase in the enyzme-substrate affinity. *MI* is maximal inhibition. *EC50* and *IC50* represent activation and inhibition constant respectively, while *p* and *q* represent the corresponding Hill’s coefficients.

## Acknowledgments

High-performance computing at the University of Rijeka is supported by the European Fund for Regional Development (ERDF) and by the Ministry of Science, Education, and Sports of the Republic of Croatia under the project number RC.2.2.06-0001. Z.M.S was a recipient of funds from the Croatian Science Foundation’s project number O-1505-2015, and for a period of time employed in medical biochemistry laboratory at the Psychiatric Hospital Rab. Z.M.S. and VM were recipients of funds from the University of Rijeka project numbers 511-12. The funders had no role in study design, data collection, and analysis, or decision to publish and preparation of the manuscript. The authors gratefully acknowledge the services of NVIDIA CUDA Teaching Center and the studies in BioSFLab (www.svedruziclab.com). Gordan Janeš, Miroslav Puškarić, and dr. Draško Tomić have provided crucial help as a part of the Center for Advanced Computing and Modeling.

We apologize that we could not include many of the relevant citations due to space limitations.

## References

1. Blennow, K., de Leon, M. J., and Zetterberg, H. (2006) Alzheimer’s disease, Lancet 368, 387–403.

2. Toyn, J. H., and Ahlijanian, M. K. (2014) Interpreting Alzheimer’s disease clinical trials in light of the effects on amyloid-β, Alzheimer’s research & therapy 6, 14.

3. Cai, T., and Tomita, T. (2020) Structure-activity relationship of presenilin in γ-secretase-mediated intramembrane cleavage, Seminars in cell & developmental biology 105, 102–109.

4. Sambamurti, K., Greig, N. H., Utsuki, T., Barnwell, E. L., Sharma, E., Mazell, C., Bhat, N. R., Kindy, M. S., Lahiri, D. K., and Pappolla, M. A. (2011) Targets for AD treatment: conflicting messages from gamma-secretase inhibitors, J Neurochem 117, 359–374.

5. Kisby, B., Jarrell, J. T., Agar, M. E., Cohen, D. S., Rosin, E. R., Cahill, C. M., Rogers, J. T., and Huang, X. (2019) Alzheimer’s Disease and Its Potential Alternative Therapeutics, Journal of Alzheimer’s disease & Parkinsonism 9.

6. Tagami, S., Yanagida, K., Kodama, T. S., Takami, M., Mizuta, N., Oyama, H., Nishitomi, K., Chiu, Y. W., Okamoto, T., Ikeuchi, T., Sakaguchi, G., Kudo, T., Matsuura, Y., Fukumori, A., Takeda, M., Ihara, Y., and Okochi, M. (2017) Semagacestat Is a Pseudo-Inhibitor of γ-Secretase, Cell reports 21, 259–273.

7. Svedružić Ž, M., Popović, K., and Šendula-Jengić, V. (2015) Decrease in catalytic capacity of γ-secretase can facilitate pathogenesis in sporadic and Familial Alzheimer’s disease, Molecular and cellular neurosciences 67, 55–65.

8. Burton, C. R., Meredith, J. E., Barten, D. M., Goldstein, M. E., Krause, C. M., Kieras, C. J., Sisk, L., Iben, L. G., Polson, C., Thompson, M. W., Lin, X. A., Corsa, J., Fiedler, T., Pierdomenico, M., Cao, Y., Roach, A. H., Cantone, J. L., Ford, M. J., Drexler, D. M., Olson, R. E., Yang, M. G., Bergstrom, C. P., McElhone, K. E., Bronson, J. J., Macor, J. E., Blat, Y., Grafstrom, R. H., Stern, A. M., Seiffert, D. A., Zaczek, R., Albright, C. F., and Toyn, J. H. (2008) The amyloid-beta rise and gamma-secretase inhibitor potency depend on the level of substrate expression, The Journal of biological chemistry 283, 22992–23003.

9. Tong, G., Wang, J. S., Sverdlov, O., Huang, S. P., Slemmon, R., Croop, R., Castaneda, L., Gu, H., Wong, O., Li, H., Berman, R. M., Smith, C., Albright, C. F., and Dockens, R. C. (2012) Multicenter, Randomized, Double-Blind, Placebo-Controlled, Single-Ascending Dose Study of the Oral gamma-Secretase Inhibitor BMS-708163 (Avagacestat): Tolerability Profile, Pharmacokinetic Parameters, and Pharmacodynamic Markers, Clin Ther 34, 654–667.

10. Svedružić, Z. M., Popovic, K., and Sendula-Jengic, V. (2013) Modulators of gamma-secretase activity can facilitate the toxic side-effects and pathogenesis of Alzheimer’s disease, PloS one 8, e50759.

11. Walsh, R. (2014) Are improper kinetic models hampering drug development?, PeerJ 2, e649.

12. Yagishita, S., Morishima-Kawashima, M., Tanimura, Y., Ishiura, S., and Ihara, Y. (2006) DAPT-induced intracellular accumulations of longer amyloid beta-proteins: further implications for the mechanism of intramembrane cleavage by gamma-secretase, Biochemistry 45, 3952–3960.

13. Gillman, K. W., Starrett, J. E., Jr., Parker, M. F., Xie, K., Bronson, J. J., Marcin, L. R., McElhone, K. E., Bergstrom, C. P., Mate, R. A., Williams, R., Meredith, J. E., Jr., Burton, C. R., Barten, D. M., Toyn, J. H., Roberts, S. B., Lentz, K. A., Houston, J. G., Zaczek, R., Albright, C. F., Decicco, C. P., Macor, J. E., and Olson, R. E. (2010) Discovery and Evaluation of BMS-708163, a Potent, Selective and Orally Bioavailable γ-Secretase Inhibitor, ACS medicinal chemistry letters 1, 120–124.

14. Jämsä, A., Belda, O., Edlund, M., and Lindström, E. (2011) BACE-1 inhibition prevents the γ-secretase inhibitor evoked Aβ rise in human neuroblastoma SH-SY5Y cells, Journal of biomedical science 18, 76.

15. Mitani, Y., Yarimizu, J., Saita, K., Uchino, H., Akashiba, H., Shitaka, Y., Ni, K., and Matsuoka, N. (2012) Differential effects between gamma-secretase inhibitors and modulators on cognitive function in amyloid precursor protein-transgenic and nontransgenic mice, J Neurosci 32, 2037–2050.

16. Fersht, A. (1998) Structure and Mechanism in Protein Science: A Guide to Enzyme Catalysis and Protein Folding (Hardcover), 1st ed., W. H. Freeman; 1st edition

17. Motulsky, H., and Christopoulos, A. (2004) Fitting Models to Biological Data Using Linear and Nonlinear Regression: A Practical Guide to Curve Fitting Oxford University Press, USA; 1 edition

18. Tipton, K. F., (Ed.) (2002) Enzyme Assays, 2nd Edition ed., Oxford University Press.

19. Svedružić Ž, M., Odorčić, I., Chang, C. H., and Svedružić, D. (2020) Substrate Channeling via a Transient Protein-Protein Complex: The case of D-Glyceraldehyde-3-Phosphate Dehydrogenase and L-Lactate Dehydrogenase, Scientific reports 10, 10404.

20. Srivastava, D. K., and Bernhard, S. A. (1987) Biophysical chemistry of metabolic reaction sequences in concentrated enzyme solution and in the cell, Annu Rev Biophys Biophys Chem 16, 175–204.

21. Svedružić, Z. M., Popovic, K., Smoljan, I., and Sendula-Jengic, V. (2012) Modulation of gamma-Secretase Activity by Multiple Enzyme-Substrate Interactions: Implications in Pathogenesis of Alzheimer’s Disease, PloS one 7, e32293.

22. Kakuda, N., Funamoto, S., Yagishita, S., Takami, M., Osawa, S., Dohmae, N., and Ihara, Y. (2006) Equimolar production of amyloid beta-protein and amyloid precursor protein intracellular domain from beta-carboxyl-terminal fragment by gamma-secretase, The Journal of biological chemistry 281, 14776–14786.

23. Hochard, A., Oumata, N., Bettayeb, K., Gloulou, O., Fant, X., Durieu, E., Buron, N., Porceddu, M., Borgne-Sanchez, A., Galons, H., Flajolet, M., and Meijer, L. (2013) Aftins increase amyloid-beta42, lower amyloid-beta38, and do not alter amyloid-beta40 extracellular production in vitro: toward a chemical model of Alzheimer’s disease?, J Alzheimers Dis 35, 107–120.

24. Chavez-Gutierrez, L., Bammens, L., Benilova, I., Vandersteen, A., Benurwar, M., Borgers, M., Lismont, S., Zhou, L., Van Cleynenbreugel, S., Esselmann, H., Wiltfang, J., Serneels, L., Karran, E., Gijsen, H., Schymkowitz, J., Rousseau, F., Broersen, K., and De Strooper, B. (2012) The mechanism of gamma-Secretase dysfunction in familial Alzheimer disease, The EMBO journal 31, 2261–2274.

25. Yagishita, S., Morishima-Kawashima, M., Ishiura, S., and Ihara, Y. (2008) Abeta46 is processed to Abeta40 and Abeta43, but not to Abeta42, in the low density membrane domains, The Journal of biological chemistry 283, 733–738.

26. Yin, Y. I., Bassit, B., Zhu, L., Yang, X., Wang, C., and Li, Y. M. (2007) {gamma}-Secretase Substrate Concentration Modulates the Abeta42/Abeta40 Ratio: implications for Alzheimer’s disease, The Journal of biological chemistry 282, 23639–23644.

27. Coric, V., van Dyck, C. H., Salloway, S., Andreasen, N., Brody, M., Richter, R. W., Soininen, H., Thein, S., Shiovitz, T., Pilcher, G., Colby, S., Rollin, L., Dockens, R., Pachai, C., Portelius, E., Andreasson, U., Blennow, K., Soares, H., Albright, C., Feldman, H. H., and Berman, R. M. (2012) Safety and Tolerability of the gamma-Secretase Inhibitor Avagacestat in a Phase 2 Study of Mild to Moderate Alzheimer Disease, Arch Neurol, 1–12.

28. Tamayev, R., and D’Adamio, L. (2012) Inhibition of gamma-secretase worsens memory deficits in a genetically congruous mouse model of Danish dementia, Mol Neurodegener 7, 19.

29. Doody, R. S., Raman, R., Farlow, M., Iwatsubo, T., Vellas, B., Joffe, S., Kieburtz, K., He, F., Sun, X., Thomas, R. G., Aisen, P. S., Siemers, E., Sethuraman, G., and Mohs, R. (2013) A phase 3 trial of semagacestat for treatment of Alzheimer’s disease, N Engl J Med 369, 341–350.

30. Henley, D. B., May, P. C., Dean, R. A., and Siemers, E. R. (2009) Development of semagacestat (LY450139), a functional gamma-secretase inhibitor, for the treatment of Alzheimer’s disease, Expert Opin Pharmacother 10, 1657–1664.

31. Bai, X. C., Rajendra, E., Yang, G., Shi, Y., and Scheres, S. H. (2015) Sampling the conformational space of the catalytic subunit of human γ-secretase, eLife 4.

32. Barnwell, E., Padmaraju, V., Baranello, R., Pacheco-Quinto, J., Crosson, C., Ablonczy, Z., Eckman, E., Eckman, C. B., Ramakrishnan, V., Greig, N. H., Pappolla, M. A., and Sambamurti, K. (2014) Evidence of a Novel Mechanism for Partial gamma-Secretase Inhibition Induced Paradoxical Increase in Secreted Amyloid beta Protein, PloS one 9, e91531.

33. Morohashi, Y., Kan, T., Tominari, Y., Fuwa, H., Okamura, Y., Watanabe, N., Sato, C., Natsugari, H., Fukuyama, T., Iwatsubo, T., and Tomita, T. (2006) C-terminal fragment of presenilin is the molecular target of a dipeptidic gamma-secretase-specific inhibitor DAPT (N-[N-(3,5-difluorophenacetyl)-L-alanyl]-S-phenylglycine t-butyl ester), The Journal of biological chemistry 281, 14670–14676.

34. Lanz, T. A., Hosley, J. D., Adams, W. J., and Merchant, K. M. (2004) Studies of Abeta pharmacodynamics in the brain, cerebrospinal fluid, and plasma in young (plaque-free) Tg2576 mice using the gamma-secretase inhibitor N2-[(2S)-2-(3,5-difluorophenyl)-2-hydroxyethanoyl]-N1-[(7S)-5-methyl-6-oxo-6,7-di hydro-5H-dibenzo[b,d]azepin-7-yl]-L-alaninamide (LY-411575), J Pharmacol Exp Ther 309, 49–55.

35. Arnarez, C., Uusitalo, J. J., Masman, M. F., Ingólfsson, H. I., de Jong, D. H., Melo, M. N., Periole, X., de Vries, A. H., and Marrink, S. J. (2015) Dry Martini, a coarse-grained force field for lipid membrane simulations with implicit solvent, Journal of chemical theory and computation 11, 260–275.

36. Zhou, R., Yang, G., Guo, X., Zhou, Q., Lei, J., and Shi, Y. (2019) Recognition of the amyloid precursor protein by human γ-secretase, Science (New York, N.Y.) 363.

37. Bai, X. C., Yan, C., Yang, G., Lu, P., Ma, D., Sun, L., Zhou, R., Scheres, S. H. W., and Shi, Y. (2015) An atomic structure of human γ-secretase, Nature 525, 212–217.

38. Skjærven, L., Yao, X. Q., Scarabelli, G., and Grant, B. J. (2014) Integrating protein structural dynamics and evolutionary analysis with Bio3D, BMC bioinformatics 15, 399.

39. Grant, B. J., Skjærven, L., and Yao, X. Q. (2020) Comparative Protein Structure Analysis with Bio3D-Web, Methods in molecular biology (Clifton, N.J.) 2112, 15–28.

40. Sato, C., Takagi, S., Tomita, T., and Iwatsubo, T. (2008) The C-terminal PAL motif and transmembrane domain 9 of presenilin 1 are involved in the formation of the catalytic pore of the gamma-secretase, J Neurosci 28, 6264–6271.

41. Takagi, S., Tominaga, A., Sato, C., Tomita, T., and Iwatsubo, T. (2011) Participation of transmembrane domain 1 of presenilin 1 in the catalytic pore structure of the gamma-secretase, J Neurosci 30, 15943–15950.

42. Gertsik, N., Am Ende, C. W., Geoghegan, K. F., Nguyen, C., Mukherjee, P., Mente, S., Seneviratne, U., Johnson, D. S., and Li, Y. M. (2017) Mapping the Binding Site of BMS-708163 on γ-Secretase with Cleavable Photoprobes, Cell chemical biology 24, 3–8.

43. Pozdnyakov, N., Murrey, H. E., Crump, C. J., Pettersson, M., Ballard, T. E., Am Ende, C. W., Ahn, K., Li, Y. M., Bales, K. R., and Johnson, D. S. (2013) γ-Secretase modulator (GSM) photoaffinity probes reveal distinct allosteric binding sites on presenilin, The Journal of biological chemistry 288, 9710–9720.

44. Ebke, A., Luebbers, T., Fukumori, A., Shirotani, K., Haass, C., Baumann, K., and Steiner, H. (2011) Novel gamma-secretase enzyme modulators directly target presenilin protein, The Journal of biological chemistry 286, 37181–37186.

45. Tian, G., Ghanekar, S. V., Aharony, D., Shenvi, A. B., Jacobs, R. T., Liu, X., and Greenberg, B. D. (2003) The mechanism of gamma-secretase: multiple inhibitor binding sites for transition state analogs and small molecule inhibitors, The Journal of biological chemistry 278, 28968–28975.

46. Ruiz-Carmona, S., Alvarez-Garcia, D., Foloppe, N., Garmendia-Doval, A. B., Juhos, S., Schmidtke, P., Barril, X., Hubbard, R. E., and Morley, S. D. (2014) rDock: a fast, versatile and open source program for docking ligands to proteins and nucleic acids, PLoS computational biology 10, e1003571.

47. Beel, A. J., Barrett, P., Schnier, P. D., Hitchcock, S. A., Bagal, D., Sanders, C. R., and Jordan, J. B. (2009) Nonspecificity of binding of gamma-secretase modulators to the amyloid precursor protein, Biochemistry 48, 11837–11839.

48. Bedi, R. K., Patel, C., Mishra, V., Xiao, H., Yada, R. Y., and Bhaumik, P. (2016) Understanding the structural basis of substrate recognition by Plasmodium falciparum plasmepsin V to aid in the design of potent inhibitors, Scientific reports 6, 31420.

49. Mahanti, M., Bhakat, S., Nilsson, U. J., and Söderhjelm, P. (2016) Flap Dynamics in Aspartic Proteases: A Computational Perspective, Chemical biology & drug design 88, 159–177.

50. Tomita, T. (2009) Secretase inhibitors and modulators for Alzheimer’s disease treatment, Expert Rev Neurother 9, 661–679.

51. Jonsson, T., Atwal, J. K., Steinberg, S., Snaedal, J., Jonsson, P. V., Bjornsson, S., Stefansson, H., Sulem, P., Gudbjartsson, D., Maloney, J., Hoyte, K., Gustafson, A., Liu, Y., Lu, Y., Bhangale, T., Graham, R. R., Huttenlocher, J., Bjornsdottir, G., Andreassen, O. A., Jonsson, E. G., Palotie, A., Behrens, T. W., Magnusson, O. T., Kong, A., Thorsteinsdottir, U., Watts, R. J., and Stefansson, K. (2012) A mutation in APP protects against Alzheimer’s disease and age-related cognitive decline, Nature 488, 96–99.

52. Potter, R., Patterson, B. W., Elbert, D. L., Ovod, V., Kasten, T., Sigurdson, W., Mawuenyega, K., Blazey, T., Goate, A., Chott, R., Yarasheski, K. E., Holtzman, D. M., Morris, J. C., Benzinger, T. L., and Bateman, R. J. (2013) Increased in Vivo Amyloid-beta42 Production, Exchange, and Loss in Presenilin Mutation Carriers, Sci Transl Med 5, 189ra177.

53. Kumar-Singh, S., Theuns, J., Van Broeck, B., Pirici, D., Vennekens, K., Corsmit, E., Cruts, M., Dermaut, B., Wang, R., and Van Broeckhoven, C. (2006) Mean age-of-onset of familial alzheimer disease caused by presenilin mutations correlates with both increased Abeta42 and decreased Abeta40, Hum Mutat 27, 686–695.

54. Koch, P., Tamboli, I. Y., Mertens, J., Wunderlich, P., Ladewig, J., Stuber, K., Esselmann, H., Wiltfang, J., Brustle, O., and Walter, J. (2012) Presenilin-1 L166P mutant human pluripotent stem cell-derived neurons exhibit partial loss of gamma-secretase activity in endogenous amyloid-beta generation, Am J Pathol 180, 2404–2416.

55. Fukumoto, H., Rosene, D. L., Moss, M. B., Raju, S., Hyman, B. T., and Irizarry, M. C. (2004) Beta-secretase activity increases with aging in human, monkey, and mouse brain, Am J Pathol 164, 719–725.

56. Theuns, J., Remacle, J., Killick, R., Corsmit, E., Vennekens, K., Huylebroeck, D., Cruts, M., and Van Broeckhoven, C. (2003) Alzheimer-associated C allele of the promoter polymorphism −22C>T causes a critical neuron-specific decrease of presenilin 1 expression, Hum Mol Genet 12, 869–877.

57. Rovelet-Lecrux, A., Hannequin, D., Raux, G., Le Meur, N., Laquerriere, A., Vital, A., Dumanchin, C., Feuillette, S., Brice, A., Vercelletto, M., Dubas, F., Frebourg, T., and Campion, D. (2006) APP locus duplication causes autosomal dominant early-onset Alzheimer disease with cerebral amyloid angiopathy, Nat Genet 38, 24–26.

58. Guyant-Marechal, L., Rovelet-Lecrux, A., Goumidi, L., Cousin, E., Hannequin, D., Raux, G., Penet, C., Ricard, S., Mace, S., Amouyel, P., Deleuze, J. F., Frebourg, T., Brice, A., Lambert, J. C., and Campion, D. (2007) Variations in the APP gene promoter region and risk of Alzheimer disease, Neurology 68, 684–687.

59. Li, R., Lindholm, K., Yang, L. B., Yue, X., Citron, M., Yan, R., Beach, T., Sue, L., Sabbagh, M., Cai, H., Wong, P., Price, D., and Shen, Y. (2004) Amyloid beta peptide load is correlated with increased beta-secretase activity in sporadic Alzheimer’s disease patients, Proceedings of the National Academy of Sciences of the United States of America 101, 3632–3637.

60. Serneels, L., Van Biervliet, J., Craessaerts, K., Dejaegere, T., Horre, K., Van Houtvin, T., Esselmann, H., Paul, S., Schafer, M. K., Berezovska, O., Hyman, B. T., Sprangers, B., Sciot, R., Moons, L., Jucker, M., Yang, Z., May, P. C., Karran, E., Wiltfang, J., D’Hooge, R., and De Strooper, B. (2009) gamma-Secretase heterogeneity in the Aph1 subunit: relevance for Alzheimer’s disease, Science (New York, N.Y.) 324, 639–642.

61. Schor, N. F. (2011) What the halted phase III gamma-secretase inhibitor trial may (or may not) be telling us, Ann Neurol 69, 237–239.

62. Miletić, V., Odorčić, I., Nikolić, P., and Svedružić Ž, M. (2017) In silico design of the first DNA-independent mechanism-based inhibitor of mammalian DNA methyltransferase Dnmt1, PloS one 12, e0174410.

63. Qi, Y., Ingolfsson, H. I., Cheng, X., Lee, J., Marrink, S. J., and Im, W. (2015) CHARMM-GUI Martini Maker for Coarse-Grained Simulations with the Martini Force Field, Journal of chemical theory and computation 11, 4486–4494.

64. Lomize, M. A., Pogozheva, I. D., Joo, H., Mosberg, H. I., and Lomize, A. L. (2012) OPM database and PPM web server: resources for positioning of proteins in membranes, Nucleic acids research 40, D370–376.

65. Van Der Spoel, D., Lindahl, E., Hess, B., Groenhof, G., Mark, A. E., and Berendsen, H. J. (2005) GROMACS: fast, flexible, and free, Journal of computational chemistry 26, 1701–1718.

66. Audagnotto, M., Kengo Lorkowski, A., and Dal Peraro, M. (2018) Recruitment of the amyloid precursor protein by γ-secretase at the synaptic plasma membrane, Biochemical and biophysical research communications 498, 334–341.

67. Krzemińska, A., Moliner, V., and Świderek, K. (2016) Dynamic and Electrostatic Effects on the Reaction Catalyzed by HIV-1 Protease, Journal of the American Chemical Society 138, 16283–16298.

68. Pettersen, E. F., Goddard, T. D., Huang, C. C., Couch, G. S., Greenblatt, D. M., Meng, E. C., and Ferrin, T. E. (2004) UCSF Chimera--a visualization system for exploratory research and analysis vesrion J Comput Chem 25, 1605–1612.

69. Lee, J., Patel, D. S., Ståhle, J., Park, S. J., Kern, N. R., Kim, S., Lee, J., Cheng, X., Valvano, M. A., Holst, O., Knirel, Y. A., Qi, Y., Jo, S., Klauda, J. B., Widmalm, G., and Im, W. (2019) CHARMM-GUI Membrane Builder for Complex Biological Membrane Simulations with Glycolipids and Lipoglycans, Journal of chemical theory and computation 15, 775–786.

70. Baker, N. A., Sept, D., Joseph, S., Holst, M. J., and McCammon, J. A. (2001) Electrostatics of nanosystems: application to microtubules and the ribosome, Proceedings of the National Academy of Sciences 98, 10037–10041.

71. Li, Y. M., Lai, M. T., Xu, M., Huang, Q., DiMuzio-Mower, J., Sardana, M. K., Shi, X. P., Yin, K. C., Shafer, J. A., and Gardell, S. J. (2000) Presenilin 1 is linked with gamma-secretase activity in the detergent solubilized state, Proceedings of the National Academy of Sciences of the United States of America 97, 6138–6143.

72. da Silva, A. W. S., and Vranken, W. F. (2012) ACPYPE-Antechamber python parser interface, BMC research notes 5, 367.

73. Kim, S., Lee, J., Jo, S., Brooks, C. L., 3rd, Lee, H. S., and Im, W. (2017) CHARMM-GUI ligand reader and modeler for CHARMM force field generation of small molecules, J Comput Chem 38, 1879–1886.

74. Lee, J., Cheng, X., Swails, J. M., Yeom, M. S., Eastman, P. K., Lemkul, J. A., Wei, S., Buckner, J., Jeong, J. C., Qi, Y., Jo, S., Pande, V. S., Case, D. A., Brooks, C. L., 3rd, MacKerell, A. D., Jr., Klauda, J. B., and Im, W. (2016) CHARMM-GUI Input Generator for NAMD, GROMACS, AMBER, OpenMM, and CHARMM/OpenMM Simulations Using the CHARMM36 Additive Force Field, Journal of chemical theory and computation 12, 405–413.

75. Humphrey, W., Dalke, A., and Schulten, K. (1996) VMD: visual molecular dynamics, Journal of molecular graphics 14, 33–38.

