## supplemental_information for "Structural Analysis of Simultaneous Activation and Inhibition of γ-Secretase Activity in Development of Drugs for Alzheimer’s disease"

**Supplemental video 1. All-atom molecular dynamic studies of penetration of DAPT molecules into the active site tunnel.** The presenilin structure is shown as a transparent white surface to show structures underneath the surface. The active site Asp 257 and Asp 358 are shown as red licorice, while the PAL motif (a.a. 433 to 435) is shown as black licorice. DAPT molecules are shown as green licorice. The Aph1 structure is shown as a transparent yellow surface. The boundaries of the lipid bilayer are illustrated by marking the water molecules with red dots. The simulation shows 200 nanoseconds of molecular events, one frame recorded every nanosecond. All four biphasic-drugs can penetrate in the active site tunnel to different depth with different rates (supp. Fig 3). We show penetration by DAPT to highlight how its flexible, elongated structure, with aromatic rings on each can adapt to dynamic changes in protein structure.

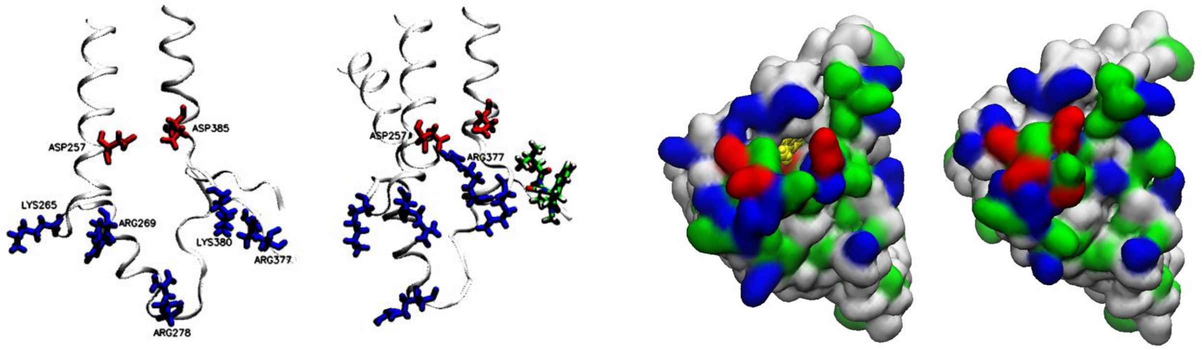

A B

**Supplement Fig. 1 (A-B). Examples of dynamic protein structures at the cytosolic end of the active site tunnel.** Molecular dynamic calculations showed that the cytosolic end of the active site tunnel is rich in charged amino acids that can form internal salt-bridges, which compete for interactions with the water molecules. (**A**) Amino acids 240 to 394 are shown as a ribbon model, positively charged amino acids are shown as blue licorice, and active site aspartates as red licorice. The figure is used to illustrates 10.7 Å motion of positive Arg 377 towards the negative Asp 257 in the active site. The salt bridges with Arg 377 can affect the pKa for the active site aspartates [1,2]. In similar motion, Arg 377 can form interactions with the negatively charged C-terminal on the nascent Aβ substrate, or with the negatively charged 3-peptide by-products (Fig. 3B). Such interactions can affect processive cleavages of the nascent Aβ catalytic intermediates. The semagacestat molecule is shown as a green licorice model to illustrate that the drugs can affect changes that control processive catalysis or regulate the opening of the active site tunnel. (**B**) The cytosolic end of presenilin structure is shown in surface mode, with amino acids colored based on charge and polarity: blue-positive, red-negative, green-polar not charged, and white-hydrophobic. The yellow VdW model illustrates the semagacestat molecule bound inside the active site tunnel in the presence of Aβ46 (Fig 7). The left model shows the structures with the tunnel open, while the right model shows the same structures after the tunnel closure.

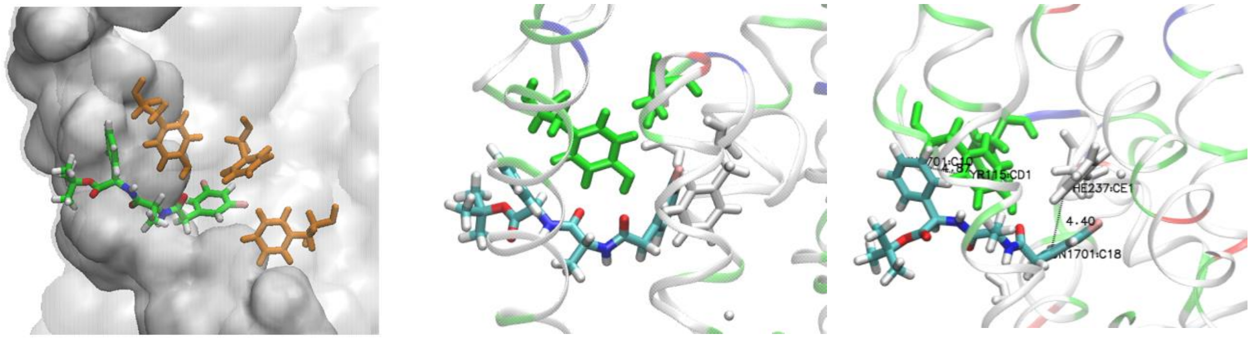

A B C

**Supplement Fig 2 (A-C) Binding interactions for DAPT molecule at the membrane-embedded end of the active site tunnel with (A) and without the substrate (B-C). (<u>A</u>)** The presenilin structure is shown as a gray transparent surface, while the substrate is shown as a gray opaque surface. DAPT molecule is shown in green, while the surrounding amino acids are shown as orange models. The substrate can block $\pi$-$\pi$ stacking interactions between the drugs and the aromatic amino acids. (**<u>B</u>-<u>C</u>**) DAPT molecule in the absence of the substrate can penetrate in the position of the substrate where it can form $\pi$-$\pi$ stacking interactions with the aromatic amino acids, most notably Phe 235 and Tyr 115.

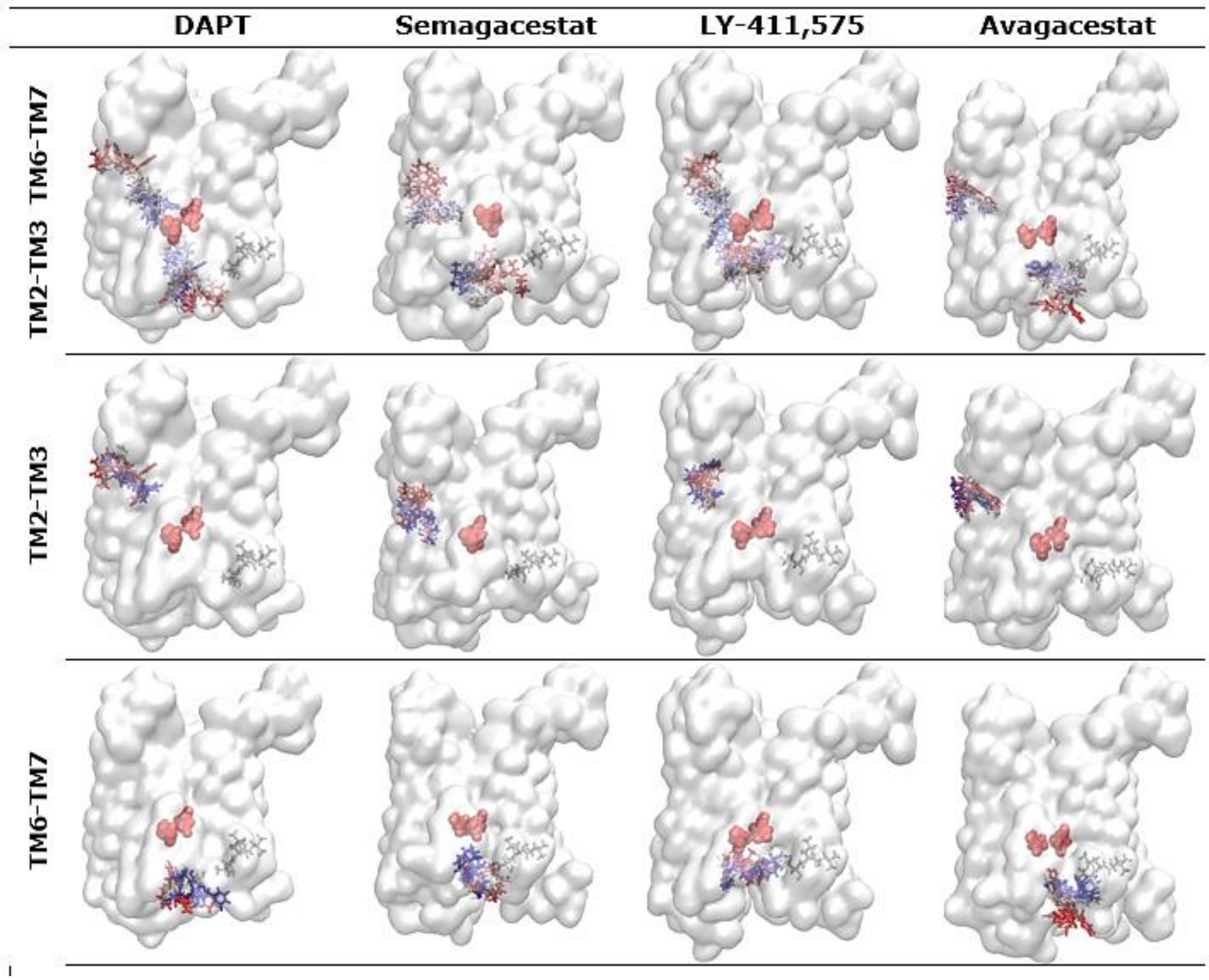

**Supplement Fig. 3. Drugs can cooperate in penetration into the active site tunnel on presenilin 1.** γ-Secretase structures with drugs bound at each end of the active site tunnel (TM2-TM3 and TM6-TM7) have been compared with the structures that had drugs bound only to one of the two ends of the tunnel (TM2-TM3 or TM6-TM7). The picture shows presenilin subunits as transparent surface, active site Asp257 and Asp385 as red VdW models, and PAL as gray licorice [3]. The binding trajectories that represent 150 nanoseconds of molecular events are depicted showing drug structures at 0, 30, 60, 90, 120, and 150 nanoseconds colored in RWB scale respectively. The starting structures for molecular dynamics calculations have been prepared using different drugs and molecular docking with γ-secretase structures with the active site tunnel closed.

1 Baker, N. A., Sept, D., Joseph, S., Holst, M. J. & McCammon, J. A. Electrostatics of nanosystems: application to microtubules and the ribosome. *Proceedings of the National Academy of Sciences* **98**, 10037-10041 (2001).
2 Krzemińska, A., Moliner, V. & Świderek, K. Dynamic and Electrostatic Effects on the Reaction Catalyzed by HIV-1 Protease. *Journal of the American Chemical Society* **138**, 16283-16298, doi:10.1021/jacs.6b06856 (2016).
3 Sato, C., Takagi, S., Tomita, T. & Iwatsubo, T. The C-terminal PAL motif and transmembrane domain 9 of presenilin 1 are involved in the formation of the catalytic pore of the gamma-secretase. *J Neurosci* **28**, 6264-6271 (2008).
